# Measuring the dynamic balance of integration and segregation underlying consciousness, anesthesia, and sleep

**DOI:** 10.1101/2024.04.12.589265

**Authors:** Hyunwoo Jang, George A. Mashour, Anthony G. Hudetz, Zirui Huang

## Abstract

Consciousness requires a dynamic balance of integration and segregation in functional brain networks. An optimal integration-segregation balance depends on two key aspects of functional connectivity: global efficiency (i.e., integration) and clustering (i.e., segregation). We developed a new fMRI-based measure, termed the integration-segregation difference (ISD), which captures both aspects. We used this metric to quantify changes in brain state from conscious wakefulness to loss of responsiveness induced by the anesthetic propofol. The observed changes in ISD suggest a profound shift to segregation in both whole brain and all brain subnetworks during anesthesia. Moreover, brain networks displayed similar sequences of disintegration and subsequent reintegration during, respectively, loss and return of responsiveness. Random forest machine learning models, trained with the integration and segregation of brain networks, identified the awake vs. unresponsive states and their transitions with accuracy up to 93%. We found that metastability (i.e., the dynamic recurrence of non-equilibrium transient states) is more effectively explained by integration, while complexity (i.e., diversity and intricacy of neural activity) is more closely linked with segregation. The analysis of a sleep dataset revealed similar findings. Our results demonstrate that the integration-segregation balance is a useful index that can differentiate among various conscious and unconscious states.

## Introduction

The brain faces the task of seamlessly binding incoming sensory information, action planning, execution, and other cognitive processes into a coherent conscious experience. This necessitates harmonizing distinct functions in spatially distributed modules and the integration of these local neural activities into a cohesive whole^1,2^. Excessive integration can lead to uncontrolled synchronization exemplified by seizure^3^, while insufficient integration can hinder the formation of a unified experience^2,4^. Consciousness is thought to critically depend on the brain’s capability to balance these two extremes, ensuring adequate integration and segregation for the unity of a meaningful computational repertoire^1,2,5,6^.

The idea of integration-segregation balance was initially conceptualized as a reformulation of the binding problem^6^. Over time, this concept has developed, establishing a connection with the neural complexity^2,6^. As part of this evolution, a measurement called “phi” was introduced to quantitatively assess integrated information^7^. However, computational challenges and ongoing debates about its neurobiological significance limit the practicality of phi^8–10^. While modified versions of phi (e.g., phi*, phi-MIP, phi-max) are applicable for smaller-scale dynamical systems, they become impractical for more complex systems, including the brain^11,12^.

Empirical studies often measure integration and segregation separately^1,5,13,14^. Graph theoretical measures such as connector hubs and participation coefficient serve as proxies for integration^14–17^, while modularity, local efficiency, and system segregation are used to gauge segregation^14,16,18^. These approaches do not capture the intertwined and mutually constraining nature of integration and segregation. These highlight the need for an empirically grounded and computationally feasible measure of the integration-segregation balance.

Inspired by the well-established concept of small-worldness^19–22^, we considered that the integration-segregation balance within the brain relies on two key aspects of functional connectivity: (1) global efficiency, a measure of how efficiently information can be exchanged across the entire network; (2) global clustering, the degree to which nodes in a network tend to cluster together and form tightly interconnected groups. In waking consciousness, efficiency (related to integration) and clustering (related to segregation and modularization) are thought to be optimized to ensure adaptive and complex brain functioning^22,23^.

Herein, we propose a new network measure, which we have named the integration-segregation difference (ISD; Fig. 1A). ISD is defined as the integration subtracted by segregation, each quantified by multi-level efficiency and clustering coefficient (see Methods for definitions). We calculate the ISD on dynamic functional connectivity of functional magnetic resonance imaging (fMRI) signals and characterize the disintegration and subsequent reintegration of brain networks during anesthetic state transitions (Fig. 1B). By applying machine learning methods, we construct a model to predict consciousness state transitions from the observed changes in integration and segregation (Fig. 1C). We examine the relationships between integration-segregation balance, metastability, and complexity—features that have been implicated as significant determinants of the state of consciousness (Fig. 1D). Finally, we reproduce our key findings in natural sleep.

**Figure 1.**
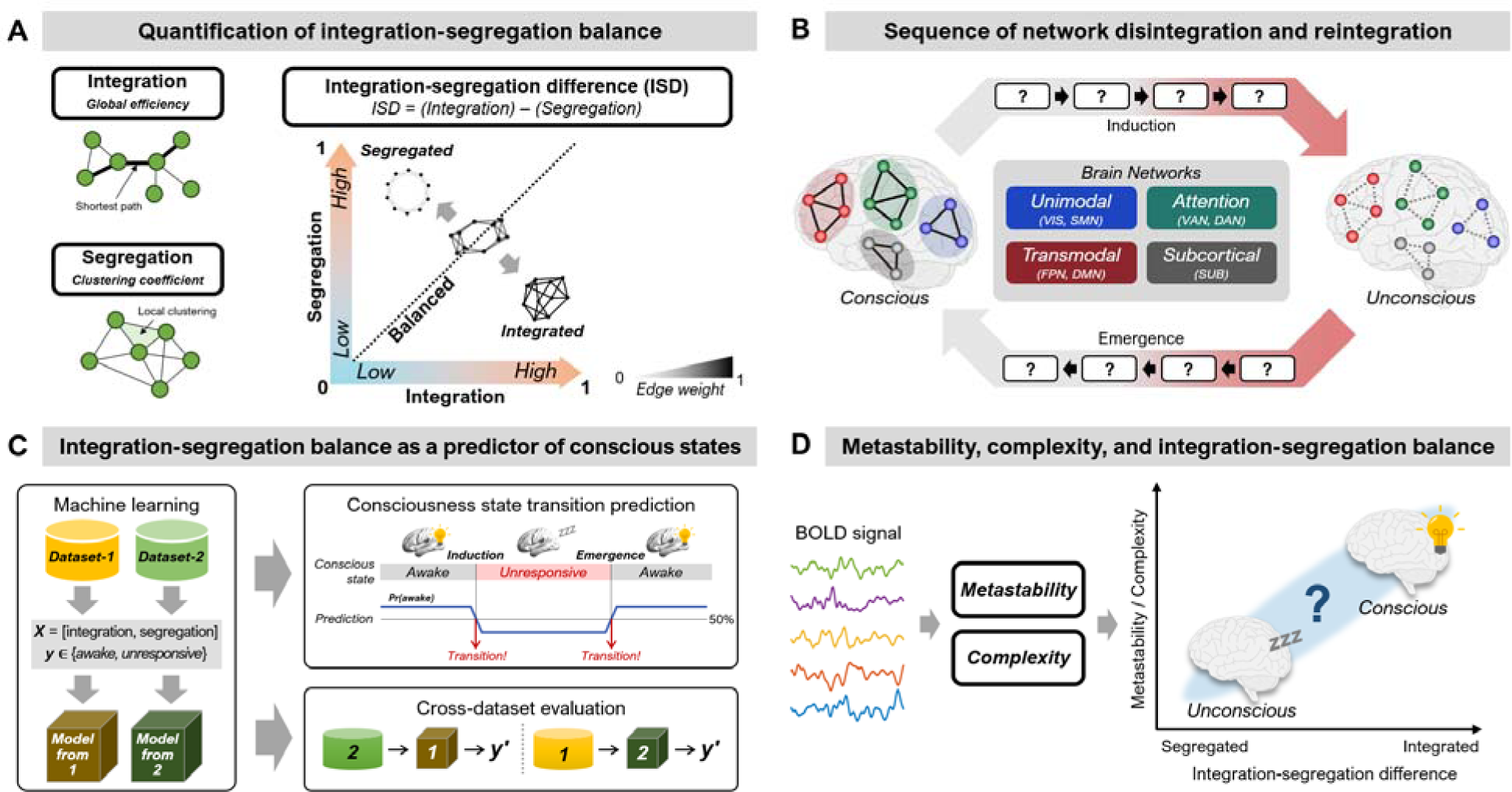
Overview of conceptual and methodological frameworks. (A) Integration-segregation balance quantified by the ISD, defined as integration (captured by efficiency) subtracted by segregation (captured by clustering). (B) The sequences of network disintegration and reintegration during induction and emergence transitions. (C) The use of integration and segregation for prediction of consciousness state transitions. The machine learning model is evaluated in a cross-dataset manner. (D) The associations between integration-segregation balance and the concepts of metastability and complexity.

## Results

We analyzed six independent fMRI datasets collected at different research sites. Datasets 1 to 4 involve intravenous infusions of propofol at various doses. Experimental conditions included conscious baseline, light and deep sedation, surgical level of anesthesia, and recovery of consciousness. Datasets 1 and 2 include the transition periods (i.e., induction and emergence). The fMRI data acquisition of Dataset-1 (*n* = 19) served as the primary data. Dataset-2 (*n* = 26) was used in combination for reproducing the key results and analyzing temporal order of network changes during transition periods, as well as for cross-dataset validation of machine learning techniques. In both Dataset-1 and Dataset-2, deep sedation was achieved by incrementally adjusting the infusion rate of propofol until behavioral responsiveness to verbal command was lost (termed loss of responsiveness; LOR). Datasets 3 and 4 that employed varied dosages of propofol, Dataset-5, which includes various sleep stages, and a Human Connectome Project dataset (Dataset-HCP) were used to validate the effectiveness of ISD index and generalizability of our findings.

### Healthy awake brain balances integration and segregation

We used Dataset-HCP (*n* = 1009) to evaluate whether healthy, awake brains maintain the balance between integration and segregation. Using our metric of integration and segregation, we observed similar values of integration (mean ± SD: 0.41 ± 0.08) and segregation (0.46 ± 0.03) (Extended Data Fig. 1) across participants. The difference between integration and segregation (ISD) was –0.05 ± 0.07, suggesting that the normal awake brain is in a balanced state, consistent with previous literature^1,17,24^.

### Integration-segregation balance shifts towards segregation during propofol-induced loss of responsiveness

During baseline, integration and segregation remained at ≈ 0.5, resulting in approximately zero ISD (Fig. 2A-C). In contrast, LOR was associated with a decrease in integration and an increase in segregation as compared to conscious baseline and recovery (Fig. 2D,E). This was associated with a statistically significant decrease in ISD values during LOR (Fig. 2F; Friedman’s ANOVA: *p* = 0.0021; Post-hoc Wilcoxon test: baseline vs. LOR *p* = 0.0050, LOR vs. recovery *p* = 0.0012; *p*-values were FDR-corrected; see Table S1 for detailed statistics), suggesting that whole-brain networks transition toward more segregated (i.e., less integrated) states under anesthesia. In recovery, ISD reverted to the baseline level. The difference between baseline and recovery was statistically not significant (Post-hoc Wilcoxon test: *p* = 0.1165). The decrease of ISD during LOR was independently replicated in Dataset-2 (Extended Data Fig. 2). Dataset-2 only showed decrease in integration, while segregation remained relatively consistent (Extended Data Fig. 2A,B). This difference, possibly caused by dose-dependency, is further described in Extended Data Fig. 3.

**Figure 2.**
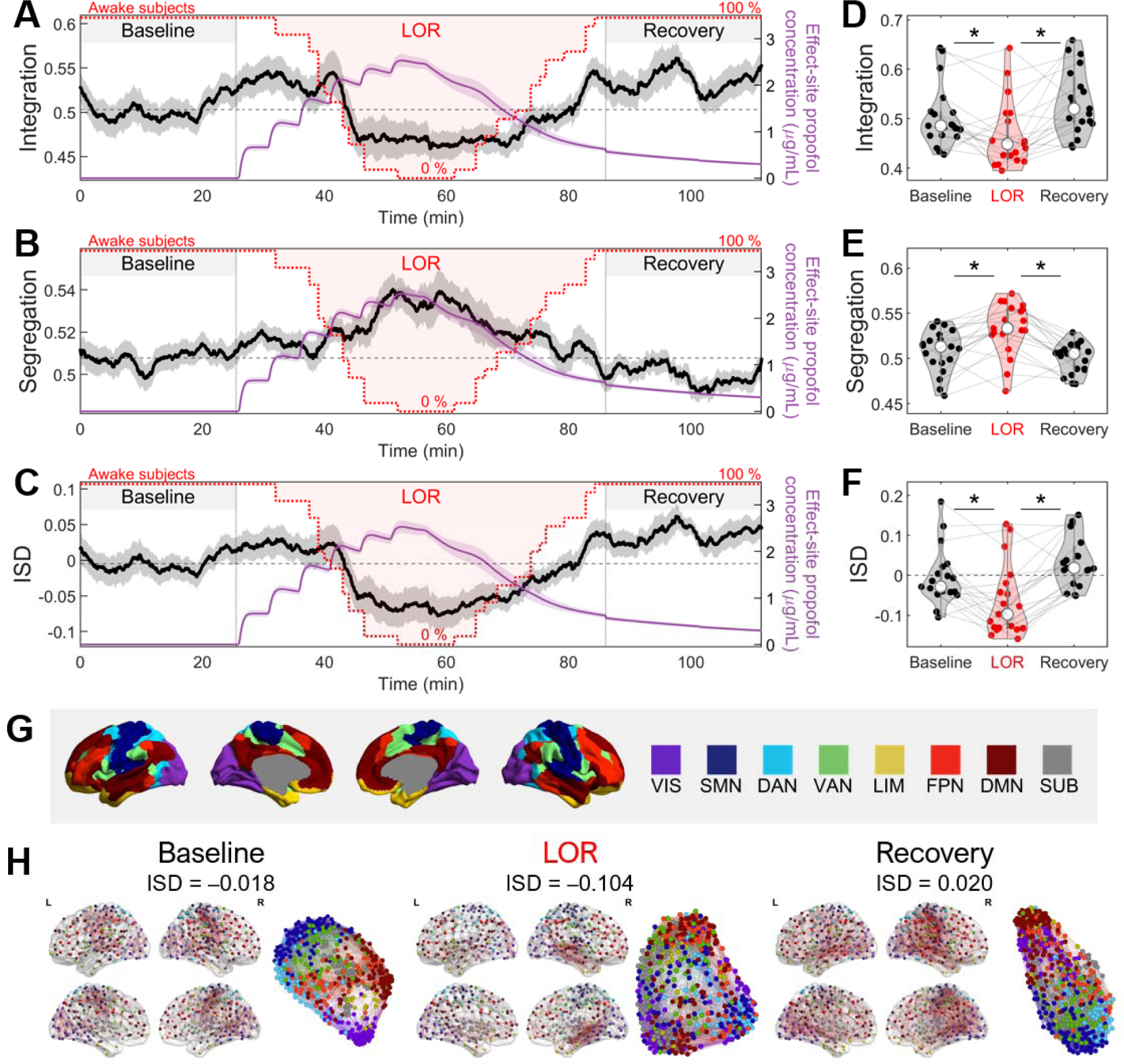
Time-resolved integration-segregation balance in the brain during propofol-induced anesthesia. (A-C) Average time course of integration, segregation, and integration-segregation difference (ISD) for Dataset-1 (*n* = 19). Horizontal dashed lines are median values during baseline. Propofol concentration is indicated by the violet line. Red line denotes the percentage of subjects responsive to verbal commands. Shaded areas represent ± SEM. Sliding window size was 4 minutes. (D-F) Average values of integration, segregation, and ISD during baseline, loss of responsiveness (LOR), and recovery states. Each point represents an individual subject, connected by grey lines. Median values are marked by white circles. Asterisks denote statistically significant differences (paired two-sided Wilcoxon signed rank test) with FDR-corrected *p* < 0.05. (G) Visualization of eight predefined functional brain networks. The networks include visual (VIS), somatomotor (SMN), dorsal attention (DAN), ventral attention (VAN), limbic (LIM), frontoparietal (FPN), default mode (DMN), and subcortical (SUB) networks. (H) Representative brain network configurations for baseline, LOR, and recovery states. Displayed are medoids of each state within the Dataset-1. Network nodes are color-coded with the assigned network.

**Figure 3.**
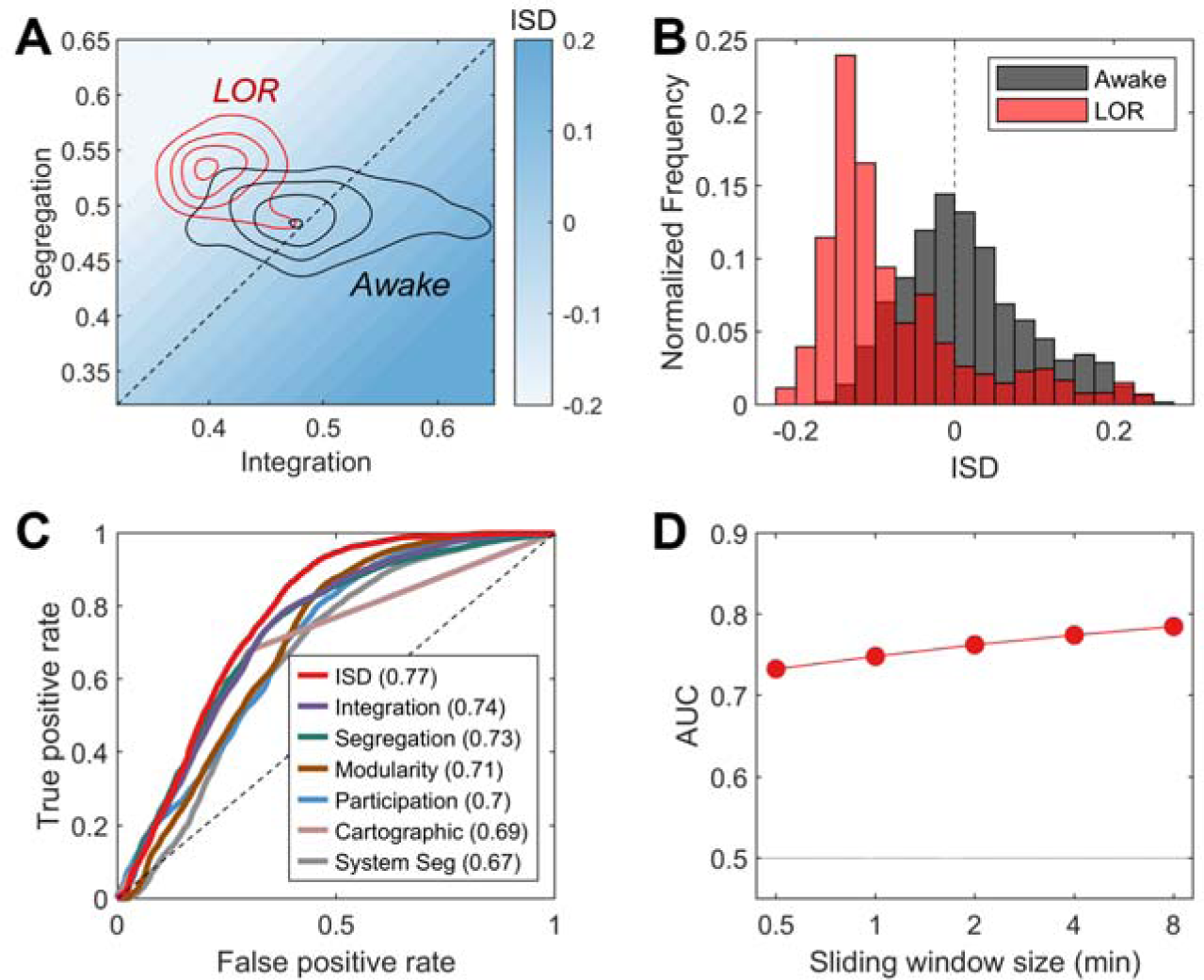
Detailed analysis of the performance of integration-segregation difference (ISD). (A) A contour density plot illustrating the distributions of awake (gray) and LOR (red) states across integration and segregation parameters. (B) Histograms of ISD value distributions for awake (gray) and LOR (red) states. (C) Receiver operating characteristics (ROC) curve analysis comparing the performance of ISD and other network metrics, quantified by area under the ROC curves (AUC). Metrics include ISD, integration (multi-level efficiency), segregation (multi-level clustering coefficient), modularity, participation coefficient, cartographic method, and system segregation. (D) The performance of ISD (as measured by AUC) in relation to the size of sliding window.

To illustrate the state-dependent changes of functional brain network architecture, we depict representative networks for each state (baseline, LOR, and recovery) (Fig. 2G,H). We selected these networks by choosing medoids of each state within the integration-segregation space (e.g., Fig. 3A) from the entire data points in Dataset-1. During baseline and recovery, networks displayed stronger connectivity. In contrast, during LOR, overall connectivity weakened, and the network became more segregated (with fewer nodes residing at the network center).

To examine the distribution of brain states in the 2D domain of integration and segregation, we used a contour density plot (Fig. 3A), which reveals distinct distributions of awake (baseline and recovery) and LOR states. In the awake state (grey), the brain resides around integration of 0.519 ± 0.077 (mean ± SD) and segregation of 0.503 ± 0.024, equivalent to balanced brain state^20,21^. In contrast, LOR state is accompanied by decrease in integration to 0.457 ± 0.073 and increase in segregation to 0.535 ± 0.037. The distributions of ISD values are markedly distinct between awake vs. LOR (Fig. 3B; awake: 0.015 ± 0.081; LOR: −0.079 ± 0.094). To assess the effectiveness of ISD in distinguishing between awake and LOR states, we evaluated its performance against other widely used metrics by comparing the area under the receiver operating characteristic curve (AUC). ISD showed superior performance in discerning awake vs.

LOR with an AUC of 0.77 (Fig. 3C), outperforming other metrics such as modularity, mean participation coefficient (the ratio of inter-network connections), cartographic method (node property profile), and system segregation (difference of intra-versus inter-network connections)^17,18,25^. The performance of ISD was robust against variations in sliding window size between 0.5 to 8 minutes (Fig. 3D). All analyses, including the superior performance of ISD, were independently replicated in Dataset-2 (Extended Data Fig. 4).

**Figure 4.**
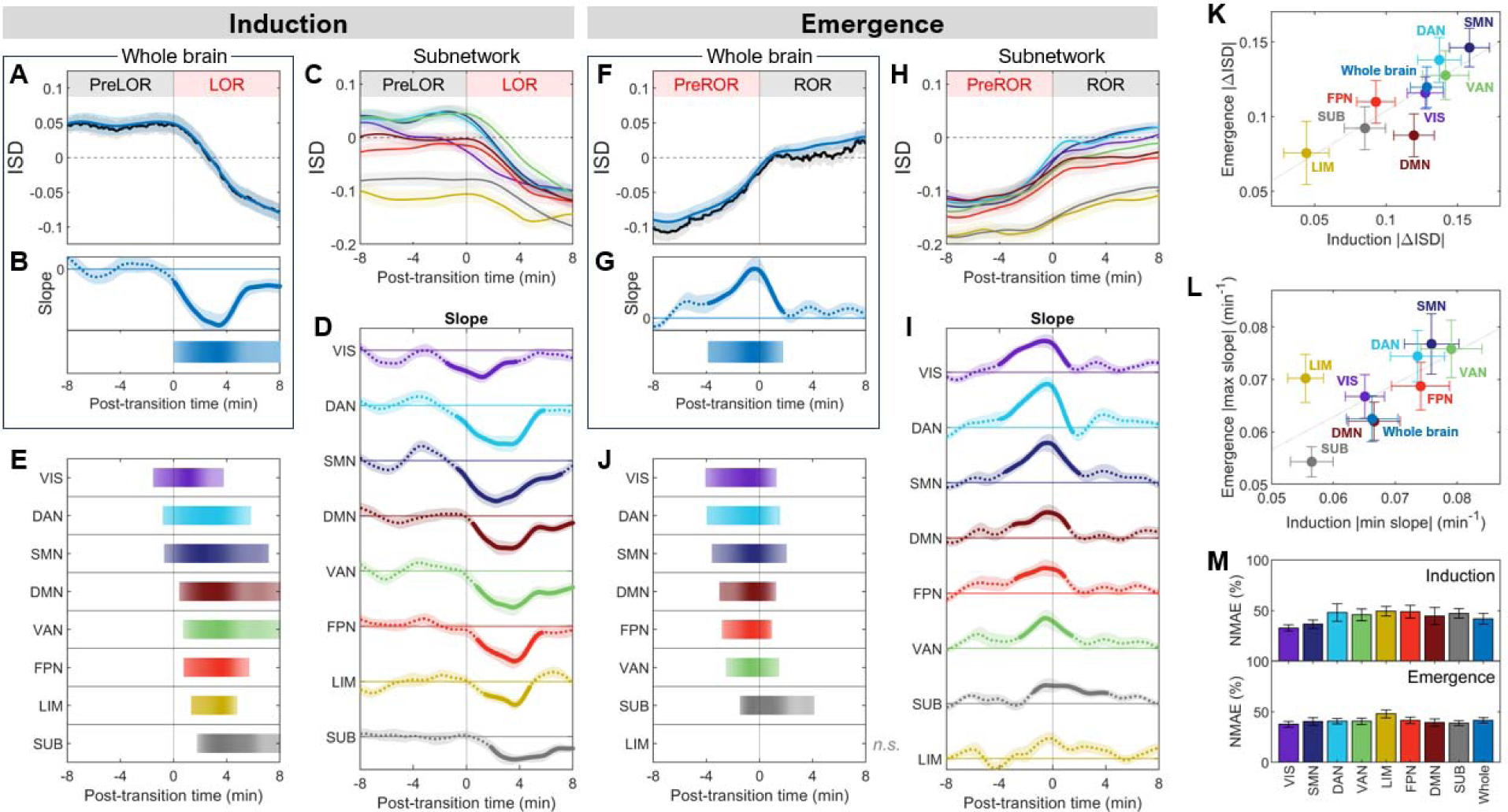
Network sequence of ISD changes during consciousness state transitions. (A, F) Group-averaged whole-brain ISD trajectories during (A) induction and (F) emergence (*n* = 45). Blue curve represent spline-smoothed averages. (B, G) (Top) Group-averaged slopes of spline-smoothed ISD trajectories. Solid regions denote periods of statistical significance (Wilcoxon sign rank test, FDR-corrected *p* < 0.05), while dashed regions indicate non-significance. (Bottom) Horizontal bars identifying the temporal regimes of statistical significance. (C, H) Smoothed averages of network ISD curves with corresponding slopes shown in (D, I). Shaded areas around the curves indicate ± SEM. (E, J) Temporal sequence of significant ISD changes across networks, sorted by the onset of significance. The *n.s.* indicates no statistical significance within the given time window. (K) Magnitude of ISD changes for induction vs. emergence. (L) Scatter plot comparing the extremes of ISD slopes during induction and emergence. (M) Bar charts of normalized mean absolute error (NMAE) between individual ISD trajectories and the group average. In all subfigures, error bars denote mean ± SEM.

To further validate the robustness and generalizability of ISD, we compared five different propofol dosage regimens (the effect-site propofol concentration [ESC] ranging from 1.0 to 4.0 *µ*g/mL) by incorporating data from two other datasets (Dataset-3 and Dataset-4; see Extended Data Fig. 3). Across all datasets, baseline conscious states (ESC = 0 *µ*g/mL) consistently stayed within specific ranges of integration (0.4 – 0.6) and segregation (0.45 – 0.55). At lower doses of propofol (ESC ≤ 2 *µ*g/mL), the brain exhibited a significant change either in integration and segregation, but not in both simultaneously (Extended Data Fig. 3C,D). For example, light sedation (ESC = 1.0 *µ*g/mL) only showed decrease in integration, whereas deep sedation (ESC = 1.9 *µ*g/mL) only exhibited increase in segregation. While integration decreased as the dose increases (except for light sedation), segregation showed non-linear trend in which it initially increases but dropped at the highest propofol dose (ESC = 4.0 *µ*g/mL), approaching a state akin to non-functional null connectivity^26^. However, by incorporating both aspects, ISD consistently captured this dose-dependent trajectory, displaying significant decreases compared to baseline across all propofol doses (Extended Data Fig. 3E).

### Temporal sequence of brain network disintegration and reintegration

We aimed to determine the timing of whole-brain ISD changes with respect to induction of and emergence from propofol-induced LOR (Fig. 4A,F). Dataset-1 and 2 were concatenated to increase statistical power (total *n* = 45). We calculated the slopes of the smoothed ISD curves from all subjects and determined the time periods when the distribution of slope values was statistically different from zero. As shown by horizontal bars in Fig. 4B, whole-brain ISD began to decrease near the onset of responsiveness loss and continued to decrease for over 8 minutes. During emergence, reintegration started earlier than recovery of responsiveness, approximately 4 minutes prior to the transition and persisted for 6 minutes (Fig. 4G).

All eight pre-defined functional brain networks underwent significant disintegration (i.e., decrease in ISD) in LOR and reintegration (i.e., increase in ISD) during recovery (Extended Data Fig. 5). Based on this, we examined if the timing of ISD changes associated with induction and emergence were distinct among the networks. Fig. 4C,D,H,I reveal network differences in the timing of ISD change for both disintegration and reintegration. During the induction phase (Fig. 4E), we observed a sequential pattern of disintegration starting from unimodal networks, such as visual (VIS) and somatomotor (SMN) networks. They began to disintegrate 1–2 minutes before LOR. This was followed by changes in the transmodal networks, such as the frontoparietal network (FPN) and default mode network (DMN). The subcortical (SUB) network was the last to disintegrate. Interestingly, the reintegration sequence observed during emergence closely resembled the order observed during induction. The reintegration began with changes in the unimodal (VIS and SMN), followed by transmodal (FPN and DMN), and finally the SUB network (Fig. 4I,J). This common sequence suggests a hierarchical pattern in the changes in network functionality.

**Figure 5.**
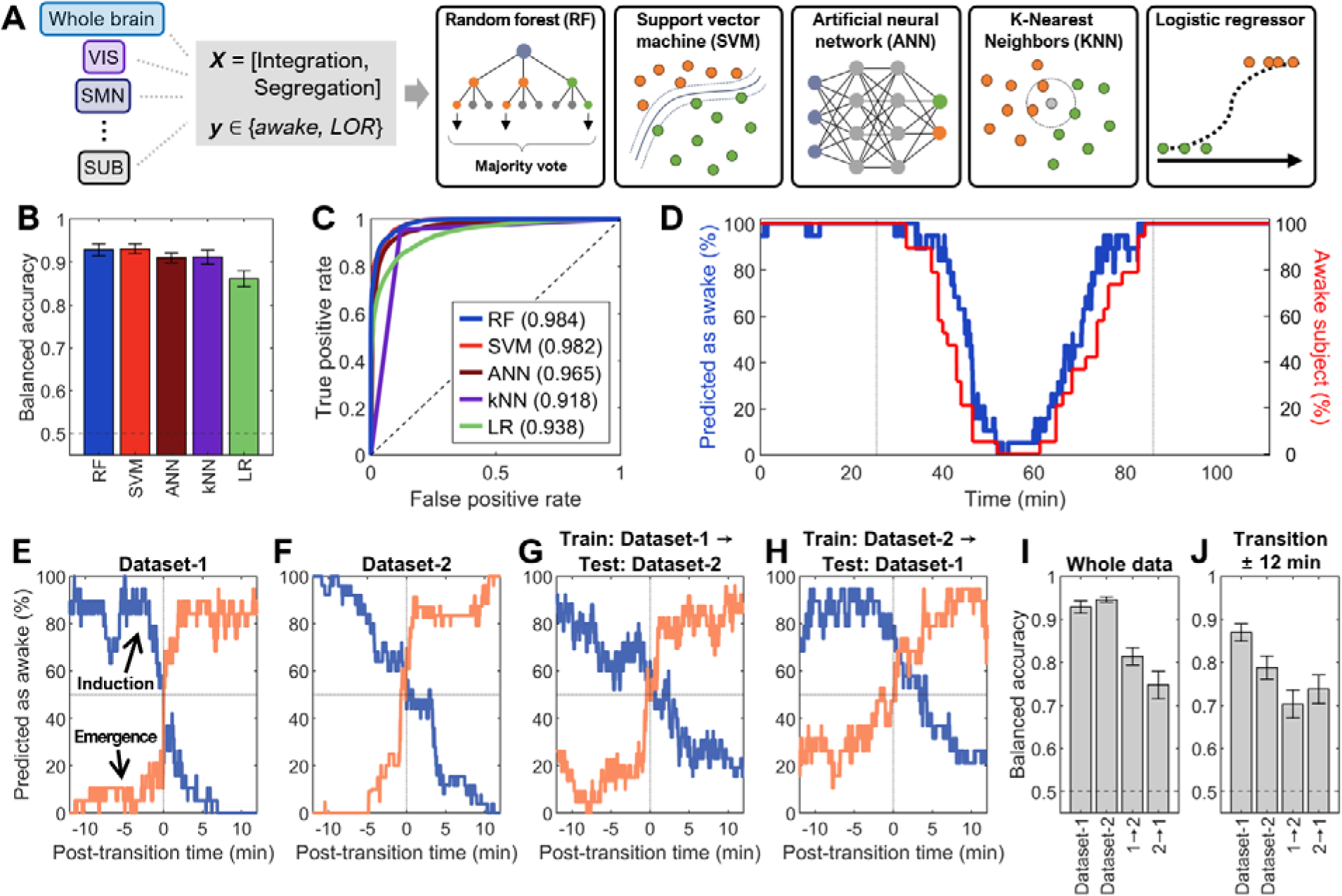
Evaluation of predictive power of integration and segregation using machine learning. (A) Schematic illustrations of supervised learning models employed: random forest (RF), support vector machine (SVM), artificial neural network (ANN), *k*-nearest neighbors (KNN), and logistic regressor (LR). (B) Comparison of model accuracies using Dataset-1 (*n* = 19). (C) ROC curves of the models (AUC values indicated in parenthesis). (D) Time course of percentage of awake state prediction by RF model (blue) vs. observed behavior states (red). (E,F) Percentage of awake state prediction by RF model around induction (blue) and emergence (orange) for (E) Dataset-1 and (F) Dataset-2. (G,H) Transferability test of the RF model trained on one dataset and tested on another dataset. (I,J) Balanced accuracy of the RF models tested across different datasets for (I) the entire data and (J) 12 minutes before and after the transitions. In all panels, error bars denote mean ± SEM across subjects during leave-one-subject-out validation.

Regarding the ISD changes (Fig. 4K), the SMN and attention networks (VAN and DAN) showed larger magnitudes of change compared to those of SUB and LIM. A similar trend was observed in the maximal slopes, i.e., the speed of change during consciousness state transitions (Fig. 4L). We also confirmed that there is a certain degree of individual variability in network transition, shown by normalized mean absolute error (NMAE) of ≈ 40% (Fig. 4M).

### Integration and segregation as predictors of states of consciousness

To evaluate if brain networks’ integration and segregation can predict the state of consciousness, we utilized a variety of supervised machine learning models (Fig. 5A). These included random forest (RF), support vector machine (SVM), artificial neural network (ANN), *k*-nearest neighbors (KNN), and logistic regressor (LR). For training, the integration and segregation values of the whole brain and all eight brain networks were given as inputs (total 18 features). Among the five models, the RF emerged as the most accurate one in predicting consciousness states, achieving an AUC of 0.984 and a balanced accuracy (average of sensitivity and specificity) of 93% when tested with Dataset-1 (Fig. 5B,C).

Predictions from the RF model closely followed the time course of behaviorally assessed states of consciousness between the induction-emergence period averaged from all participants (Fig. 5D). This alignment was particularly pronounced around the transition periods (12 minutes before and after induction or emergence; Fig. 5E). The accuracy of the RF model was high during the transition periods (balanced accuracy of 87%). We emphasize that these transition periods were not included in the training data, underscoring the model’s generalizability (Fig. 5I,J). When analyzing Dataset-2 with the RF model, we observed a similar trend as was evident in Dataset-1 (Fig. 5F,I,J). The cross-dataset performance of the RF model was evaluated by training it on one dataset and then testing it on another. These transferability tests revealed an average accuracy of approximately 75% (Fig. 5G-J).

### The link between integration-segregation balance and metastability

It has been hypothesized that integration-segregation balance may affect the degree of metastability of brain networks^3,27^. Metastability, the dynamic recurrence of non-equilibrium transient states, facilitates the brain’s fluid transition between different neural configurations and the ease of overcoming energy barriers^27^, which is vital for responding to diverse cognitive demands^3^. To quantify brain’s tendency for state switching, we utilized the moving standard deviation of the Kuramoto order parameter *R* as a proxy (Fig. 6A)^28,29^. Low metastability (Fig. 6B) is characterized by reduced dynamics between synchronized and desynchronized states, whereas high metastability (Fig. 6C) is associated with abrupt shifts between two states. Such fluctuations are illustrated by the temporal profiles of the Kuramoto order parameter (Fig. 6D).

**Figure 6.**
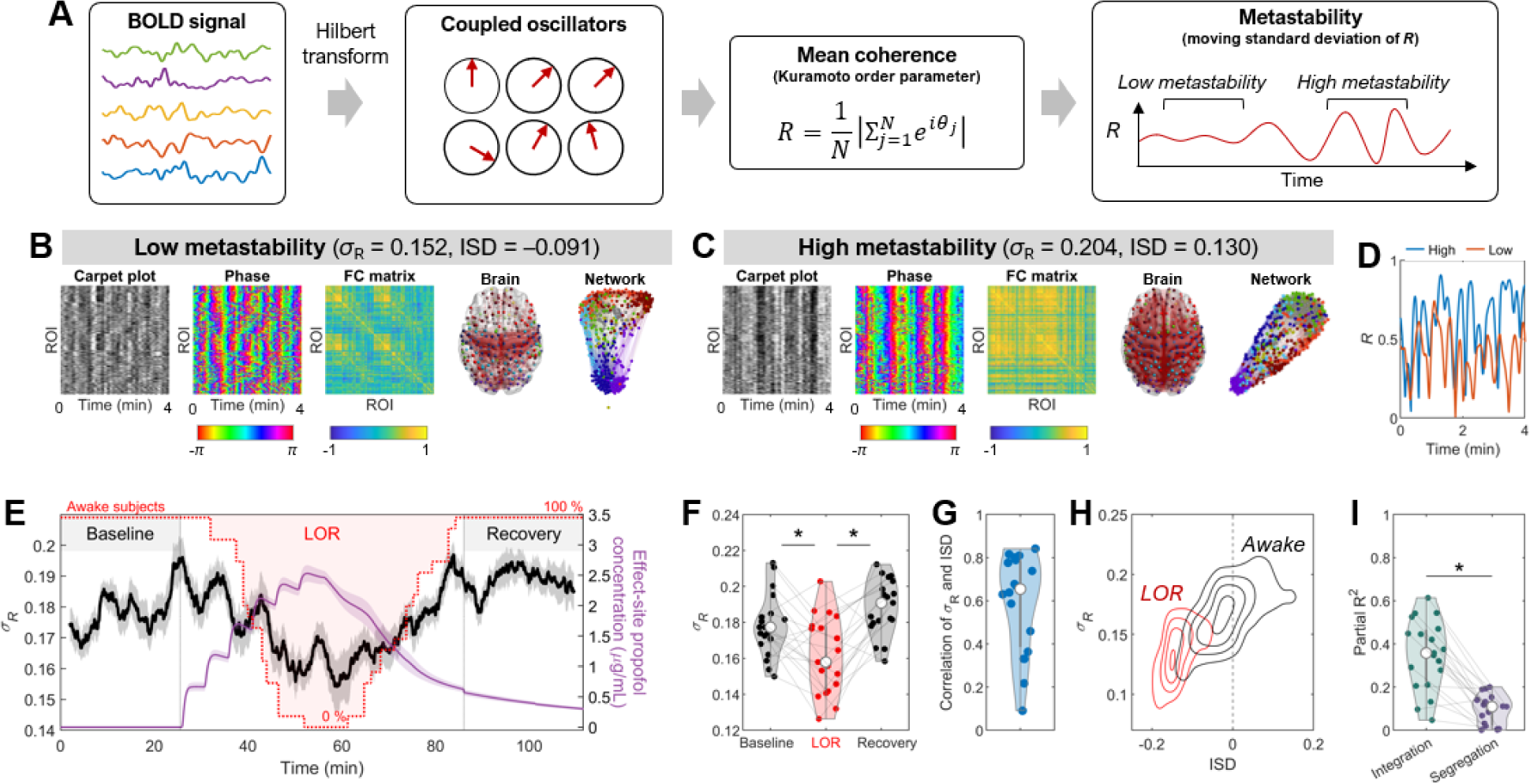
Integration-segregation balance and metastability. (A) Schematic illustration of metastability calculation process. BOLD signal is transformed to phases via the Hilbert transform. Dynamic metastability is quantified as moving standard deviation of Kuramoto order parameter (*R*). (B, C) Representative time periods with (B) low (16.7% percentile) and (C) high (83.3% percentile) metastability values, with corresponding carpet plots, functional connectivity (FC) matrices, brain graphs, and network plots. (D) Temporal variation of the Kuramoto order parameter *R*, corresponding to the time periods shown in (B) and (C). (E) A time course of metastability during loss and recovery of responsiveness in Dataset-1. (F) State-averaged values of metastability. (G) Spearman correlation of metastability and ISD measured for individual subjects. (H) A contour density plot to distinguish awake and LOR states in terms of ISD and metastability. (I) Partial *R*^2^ from a dominance analysis relating metastability to integration and segregation. The asterisk denotes statistically significant differences (paired two-sided Wilcoxon signed rank test) with *p* < 0.05.

The time course (Fig. 6E) and state-averaged distributions (Fig. 6F) of metastability tracked the changes of consciousness states with a significant reduction during LOR as compared to baseline and recovery (Friedman’s ANOVA: *p* = 0.0004; Post-hoc Wilcoxon test: baseline vs. LOR, *p* = 0.0134; LOR vs. recovery, *p* = 0.0016; baseline vs. recovery, *p* = 0.1075; *p*-values were FDR-corrected; see Table S2 for detailed statistics). We also found that metastability is highly correlated with ISD (Spearman correlation: *ρ* = 0.591 ± 0.226; Fig. 6G,H) To evaluate the individual influences of integration and segregation on metastability, we applied a dominance analysis (Fig. 6I)^30,31^. Integration explained more variance of metastability than segregation did (Partial *R*^2^ = 0.263 ± 0.190 for integration; Partial *R*^2^ = 0.054 ± 0.040 for segregation; *p* = 1.62 × 10^-4^), underscoring the relative importance of integration in explaining the brain’s metastable phenomenon. These findings were independently replicated in Dataset-2 (Extended Data Fig. 6A-C).

### The link between integration-segregation balance and complexity

We also investigated how the complexity of fMRI signal relates to integration-segregation balance in different states of consciousness^32^. Optimal complexity is thought to help the brain attain the specialized functions of segregated modules, while maintaining a sufficient level of integration^33^. By applying temporal *k*-means clustering to fMRI signal, we converted the BOLD time series into a categorical sequence of patterns (Fig. 7A)^34^. We measured three distinct complexity proxies: entropy of pattern occurrence, entropy of transition matrix, and information-theoretical complexity measure effort-to-compress^33,35^. We observed high correlations among these proxies (Extended Data Fig. 7). For simplicity, we linearly combined them into one unified metric (termed pattern complexity; see Methods for more details). This allowed us to assess both the diversity and temporal evolution in brain activity patterns. The period with low pattern complexity was marked by more regular and lattice-like brain state with weaker connectivity (Fig. 7B) than the high pattern complexity period (Fig. 7C).

**Figure 7.**
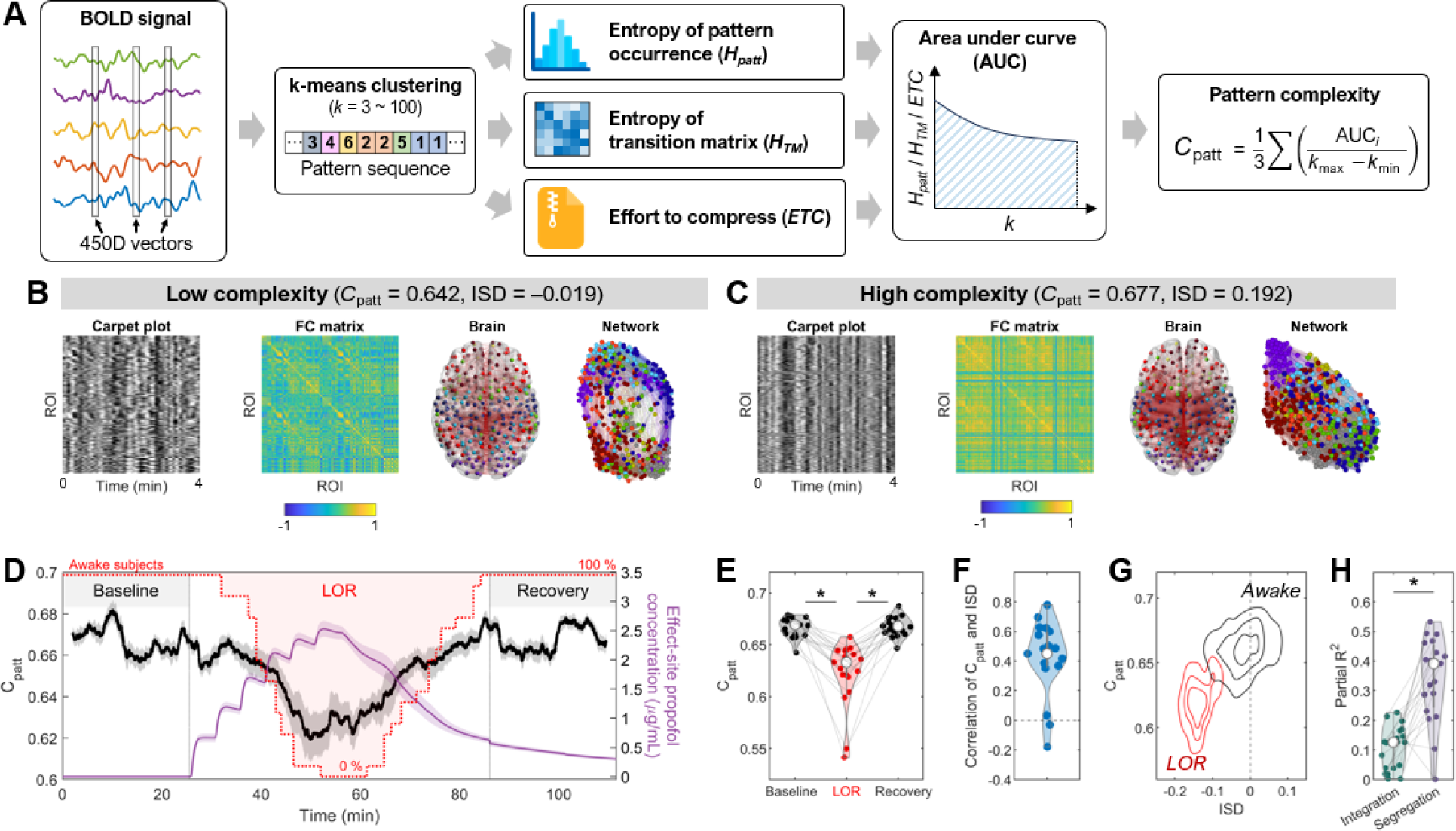
Integration-segregation balance and pattern complexity. (A) Schematic illustration of pattern complexity calculation process. BOLD signals are converted to pattern sequence via *k*-means clustering (*k* = 3 – 100). The entropy values of pattern occurrence (*H_patt_*) and transition matrix (*H_TM_*), a well as effort-to-compress (*ETC*), are calculated for each *k*. Pattern complexity is defined as the mean of the normalized areas under the curve (AUC) of *H*_patt_, *H*_TM_, and *ETC*. (B, C) Representative time period with (B) low (16.7% percentile) and (C) high (83.3% percentile) pattern complexity values, with corresponding carpet plots, functional connectivity (FC) matrices, brain graphs, and network plots. (D) A time course of pattern complexity. (E) State-averaged values of pattern complexity. (F) Spearman correlation of pattern complexity and ISD measured for individual subjects. (G) A contour density plot distinguishing between awake and LOR states in terms of ISD and pattern complexity. (H) Partial *R*^2^ from a dominance analysis relating complexity to integration and segregation. Asterisks denote statistically significant differences (paired two-sided Wilcoxon signed rank test) with *p* < 0.05.

The time course (Fig. 7D) and state-average distributions (Fig. 7E) of pattern complexity also closely followed the shifts in states of consciousness (Friedman’s ANOVA: *p* = 5.75 × 10^-7^; Post-hoc Wilcoxon test: baseline vs. LOR, *p* = 0.0002; LOR vs. recovery, *p* = 0.0002; baseline vs. recovery, *p* = 0.8092; *p*-values were FDR-corrected; see Table S3 for detailed statistics). We found that complexity is highly correlated with ISD (Spearman correlation: *ρ* = 0.419 ± 0.285) (Fig. 7F,G) The dominance analysis revealed that segregation explained larger amount of variance in complexity (Partial *R*^2^ = 0.292 ± 0.180) than integration did (Partial *R*^2^ = 0.042 ± 0.154; *p* = 0.0004) (Fig. 7H). These findings were independently replicated in Dataset-2 (Extended Data Fig. 6D-F).

### Integration-segregation balance shifts towards segregation during natural sleep

We next aimed to determine whether our findings during pharmacological states of unconsciousness induced by propofol extend to physiological states of unconsciousness during natural sleep. We analyzed an fMRI dataset acquired during various stages of sleep in humans (Dataset-5)^36,37^. Integration showed statistically significant decrease during N2 sleep compared to wakefulness and N1 sleep (Fig. 8A; Friedman’s ANOVA: *p* = 0.0058; Post-hoc Wilcoxon test: awake vs. N2, *p* = 0.0322; N1 vs. N2, *p* = 0.0322; awake vs. N1, *p* = 0.1329; *p*-values were FDR-corrected). Segregation did not show a statistically significant change (Fig. 8B; Friedman’s ANOVA: *p* = 0.2484). Driven by the changes in integration, ISD significantly decreased during N2 sleep compared to wakefulness and N1 sleep (Fig. 8C; Friedman’s ANOVA: *p* = 0.0087; Post-hoc Wilcoxon test: awake vs. N2, *p* = 0.0222; N1 vs. N2, *p* = 0.0222; awake vs. N1, *p* = 0.1943; *p*-values were FDR-corrected).

**Figure 8.**
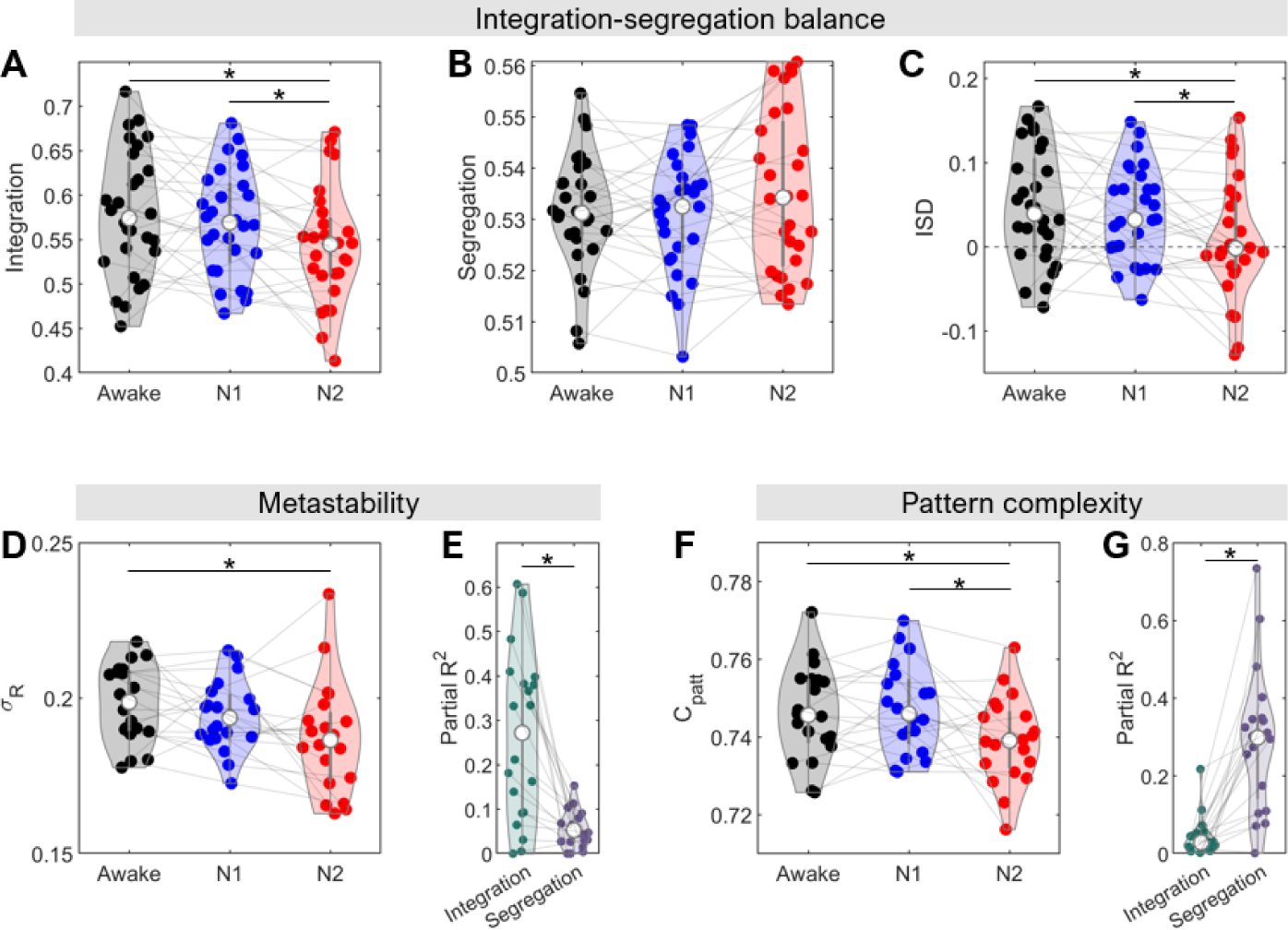
Changes in integration-segregation balance, metastability, and pattern complexity during natural sleep. (A) Integration, (B) segregation, (C) ISD, (D) metastability, and (F) pattern complexit during wakefulness and various sleep stages (*n* = 28 for integration-segregation balance analysis; *n* = 20 for metastability and pattern complexity analysis). (E) Partial *R*^2^ from a dominance analysis relating metastability to integration and segregation. (G) Partial *R*^2^ from a dominance analysis relating pattern complexity to integration and segregation. Asterisks denote statistically significant differences (paired two-sided Wilcoxon signed rank test) with FDR-corrected *p* < 0.05.

Metastability decreased at N2 sleep compared to wakefulness and N1 sleep (Fig. 8D; Friedman’s ANOVA: *p* = 0.0078; Post-hoc Wilcoxon test: awake vs. N2, *p* = 0.0334; N1 vs. N2, *p* = 0.0657; awake vs. N1, *p* = 0.1560; *p*-values were FDR-corrected). Pattern complexity also showed a decrease during N2 sleep (Fig. 8F; Friedman’s ANOVA: *p* = 0.0407; Post-hoc Wilcoxon test: awake vs. N2, *p* = 0.0414; N1 vs. N2, *p* = 0.0414; awake vs. N1, *p* = 0.6274; *p*-values were FDR-corrected). Notably, the relationships between integration, segregation, metastability, and complexity observed under anesthesia (Fig. 6,7) were consistently found in the sleep data. Metastability was more explained by integration (Partial *R*^2^ = 0.263 ± 0.054) than segregation (Partial *R*^2^ = 0.189 ± 0.040; *p* = 0.0002; Fig. 8E), while pattern complexity was more explained by segregation (Partial *R*^2^ = 0.292 ± 0.180) than integration (Partial *R*^2^ = 0.041 ± 0.049; *p* = 0.0001; Fig. 8G). See Table S4 for detailed statistics.

### Control analyses accounting for potential head motion effect

We investigated whether variations in ISD, metastability, and pattern complexity were caused by head motion differences across consciousness states. To control head motion effects, we conducted ANOVA and post-hoc tests with frame-wise displacement (FD) values regressed out. We found head motion had negligible impact (Table S1-3), showing that the observed changes in ISD, metastability, and pattern complexity are not attributable to motion artifacts.

## Discussion

We demonstrated that a novel measure, integration-segregation difference (ISD), could track changes in states of consciousness during the induction, maintenance, and emergence phases of propofol-induced loss of responsiveness (LOR). Moreover, we uncovered a common temporal sequence of network changes that occur during the transition into and out of anesthetic-induced LOR. Employing machine learning models, we showed that ISD can predict the time course of changes in states of consciousness. Further analysis revealed that metastability and complexity depend on distinct aspects of integration-segregation balance in the brain. Specifically, metastability is linked to network integration, while complexity is more associated with network segregation. These results were generalizable to natural sleep, which suggest that the integration-segregation difference may be useful to objectively identify changes in states of consciousness, with potential implications to assess disorders of consciousness.

Consciousness hinges on the brain’s ability to simultaneously integrate information to foster unity and segregate subprocesses to support specialized computations^7^. This requires an optimal balance of integration and segregation, which depend on functional and topological properties of brain networks^1^. Prior studies have typically measured either integration and segregation independently^5,18^ or focused solely on one of these aspects^24,38^. In commonly-used cartographic methods, brain states are clustered to either predominantly integrated or segregated states based on node property distribution^25^. This method faces limitations due to its binary nature which hinders a nuanced quantification of a balanced state. Also, since it performs binary clustering with dynamic functional connectivity on an individual level (i.e., each individual’s centroids may differ), it is suboptimal for group-level generalization.

The integration-segregation difference (ISD) is proposed as a method to overcome these limitations. To address the inherent weighted nature of edge distribution in functional connectivity matrix, we binarized the functional connectivity matrix at different thresholds (between 0 and 1) and calculated the global efficiency and clustering coefficient thereof. This multi-level method enables us to account for network properties in various topological scales or levels^39^. Network integration and segregation were defined as the average efficiency and clustering values across the threshold range. We then subtracted segregation from integration to obtain ISD, a single metric of integration-segregation balance. As a continuous variable, ISD can quantify the dynamics of brain states and the associated changes in consciousness, thereby extending former approaches.

Exploring the sequence of brain network transitions during changes of consciousness offers insights into the mechanism(s) of dynamic dissolution of brain functions leading to unconsciousness as well as their subsequent reanimation as consciousness is regained. Although several studies have examined the sequence of network transitions during sleep^40,41^, the body of research in the context of anesthetic-induced loss of consciousness has been limited^42^. Our present findings illustrate a similar ordered sequence of disintegration and reintegration, aligning with the induction and emergence phases of anesthesia. During induction, the earlier disintegration of unimodal networks than that of higher-order transmodal networks (e.g., default-mode and frontoparietal networks) might reflect a cascading reduction of the brain’s ability to synthesize sensory information. While sensory processing is suppressed, frontal-executive functions may be partially maintained^43^. Notably, disintegration of higher-order networks begins after the loss of responsiveness, suggesting that a certain degree of intrinsic information processing still occurs in higher-order regions even during the unresponsive state^44,45^. However, the disintegration of (whole-brain and lower-order) functional networks may hinder global or network-level broadcasting of incoming and outgoing signals, leading to loss of responsiveness^4,45^.

During emergence, the sequence of reintegration followed the similar order observed during induction. Again, unimodal brain systems exhibited earlier recovery compared to transmodal systems, implying that sensory capabilities recover prior to frontal-executive functions. Also, relatively early recovery of the dorsal attention network might reflect a reboot of attention and goal-directed behavior prior to other cognitive functions^42^. A notable difference observed during the emergence process compared to induction is that the reintegration of all networks starts ≈ 4 minutes before and culminates at recovery of responsiveness. This observation aligns with findings from EEG and behavioral studies indicating that frontal-parietal dynamics return to baseline levels before the restoration of responsiveness^42^. Additionally, both the behavior responsiveness and the dynamics of integration and segregation across the whole brain exhibited hysteresis in relation to the concentration of propofol where recovery occurs (e.g., Fig. 2C). These findings lend support to the concept of neural inertia, which suggests an inherent neuronal resistance to changes in behavioral states^46,47^. Our work also aligns with previous studies that demonstrated asymmetrical neural dynamics during the loss and regain of consciousness^48,49^.

We also used machine learning to demonstrate the ability of integration and segregation to predict the state of consciousness. By employing the random forest model, we were able to forecast transitions from awake to LOR states, or vice versa, with accuracy higher than 80%. The model’s ability to generalize effectively across different datasets highlights the robustness of network integration and segregation as a predictive metric and implies clinical usefulness. For example, integration-segregation balance has potential for objectively identifying states in disorders of consciousness without a reference to behavioral indicators that are unavailable in non-communicating patients^50,51^.

Our results revealed a relationship between integration-segregation balance and metastability^27,52^. Although previous studies have explored their connections^3,27^, it has remained understudied empirically, especially in the context of altered states of consciousness. We found a strong correlation between metastability and the integration-segregation balance, mainly driven by network integration. This implies that elevated metastability may necessitate a strong level of global efficiency, or vice versa. In addition, our results support the notion that metastability could represent a neural signature of system dynamics associated with the emergence of consciousness, as proposed by complex systems theory^27,28^.

The other related quantity, complexity, indicates the diversity and richness of neural patterns^2^. Since the conception of the integration-segregation balance, its relationship with complexity and consciousness has been a topic of interest^2,6^. However, defining and quantifying complexity in brain signals, especially with the low temporal resolution of fMRI, presents challenges due to the varied and often computationally intensive definitions. This has led to a lack of standardized methods for assessing fMRI signal complexity. To address this gap, we introduced a novel metric called pattern complexity, which leverages the high spatial resolution of fMRI. This method quantifies the uniformity vs. non-triviality of the fMRI signals as the brain navigates through a high-dimensional state space. Our findings reveal a decrease in pattern complexity concurrent with the loss of consciousness, consistent with previous studies using different complexity measures^53,54^. Furthermore, our observations reveal that pattern complexity is more closely associated with segregation than with integration. This suggests that the complexity of neural dynamics relies more heavily on the localized topological architecture, which facilitates complex neural trajectories, rather than on global interconnectedness. This underscores the importance of segregated, specialized neural networks in maintaining the intricate processing capabilities essential for conscious experience.

A few limitations of our study are recognized. Although ISD decrease was commonly observed during anesthesia and sleep, expanding our research to encompass various anesthetics, psychedelic experiences, and disorders of consciousness would improve the generalizability of our findings. The observed network order during consciousness state transitions is specific to propofol-induced anesthesia. Our use of a 4-minute sliding window, although adequate for tracking longer-term changes in consciousness states during anesthesia, restricts our ability to detect transient fluctuations at shorter time scales. Future studies could consider adjusting the window length or incorporating the instantaneous phase coherence method^55^ to capture these transient fluctuations, thus offering a more complete insight.

In conclusion, we presented a new metric of the brain’s integration-segregation balance, with initial evidence that it can assess changing states of consciousness in a principled way. This approach can be broadly applied across diverse research domains, enabling thorough examinations of brain network dynamics, including their relevance to pharmacological, neuropathological, and psychiatric conditions. By linking multiple facets of brain dynamics, such as metastability and complexity, our findings provided further insights into the concept of integration-segregation balance and its relation to consciousness.

## Methods

### Dataset-1

The dataset utilized in this study has been the subject of prior publications^48,56,57^, wherein it was analyzed using methodologies distinct from those employed in the current research. The experimental protocol received approval from the Institutional Review Board (IRB) at the University of Michigan, ensuring compliance with all pertinent guidelines and regulations. This study involved twenty-six healthy individuals (gender distribution: 13 males and 13 females; age range: 19-34 years; all right-handed). Before participating, everyone provided informed consent and was compensated upon completion of their participation in the experiment. All the participants were assessed and deemed to be in the American Society of Anesthesiologists physical status 1.

Prior to the commencement of the study, all participants were required to adhere to an eight-hour fasting period. On the experiment day, a preoperative evaluation, including a physical examination, was conducted by an attending anesthesiologist. The experiment was continuously overseen by two experienced anesthesiologists who were on-site to monitor vital signs such as spontaneous breathing, end-tidal CO_2_, heart rate, pulse oximetry, and the electrocardiogram. Noninvasive arterial pressure was recorded using an MR-compatible automatic monitoring device. Participants received a local anesthetic through a subcutaneous injection of 0.5 mL of 1% lidocaine, followed by the insertion of an intravenous cannula. They were also administered supplemental oxygen at a flow rate of 2 L/min through a nasal cannula.

We used propofol as the reference sedative-hypnotic, widely recognized for its prevalent use in fMRI research to investigate anesthesia’s impacts on human subjects. Propofol is effective due to its minimal influence on cerebral blood flow and its capacity for precise titration. The drug operates by potentiation of GABA-A receptor-mediated inhibition across the brain. The administration of propofol was meticulously managed through a target-controlled intravenous bolus and a steady infusion regimen. The specific bolus dose, rate of infusion, and duration were established in advance, following the Marsh pharmacokinetic model as integrated into the STANPUMP software (http://opentci.org/code/stanpump). The dosing strategy, combining bolus and continuous infusion, was adjusted in 5-minute intervals to achieve the desired concentration at the effect site. The anesthetic depth was gradually adjusted using a stepwise increase in dosage (0.4 μg/mL increments) until the subjects exhibited no behavioral reactions^48^. The target concentration was sustained for 21.6 ± 10.2 min, after which the infusion ceased, permitting the subjects to recover consciousness naturally.

The assessment of behavioral responsiveness involved participants squeezing a rubber ball, a method used to determine the onset of deep sedation induced by propofol, characterized by the absence of behavioral response. The BIOPAC (https://www.biopac.com) MP160 system, coupled with AcqKnowledge software version 5.0, was employed to quantify these responses, measuring the exerted air pressure in mmHg. During the scan, a series of 60 motor response trials were executed, spaced at approximately 90-second intervals. Each trial commenced with the command “action,” prompting participants to squeeze the ball once. Between these motor assessments, participants engaged in various mental imagery exercises, such as envisioning playing tennis, navigating space, or imagining the act of squeezing. Further details on the experimental setup are elaborated in prior publications^48,57^.

Data collection was performed at Michigan Medicine, University of Michigan, utilizing a 3T Philips MRI scanner equipped with a standard 32-channel transmit/receive head coil. High-resolution anatomical images were captured using T1-weighted spoiled gradient recalled echo (SPGR) imaging, employing parameters of 170 sagittal slices, 1.0 mm slice thickness without interslice gaps, a repetition time (TR) of 8.1 s, an echo time (TE) of 3.7 ms, a flip angle of 8°, a field of view (FOV) of 240 mm, and an image matrix of 256 x 256.

Functional brain images were obtained using a gradient-echo echo-planar imaging (EPI) sequence, with settings of 28 slices, TR/TE = 800/25 ms by multiband acquisition (MB factor = 4), slice thickness = 4 mm, in-plane resolution = 3.4 x 3.4 mm, FOV = 220 mm, flip angle = 76°, and a 64 x 64 image matrix. Before the upgrade of MRI hardware, six subjects underwent scans with slightly different parameters: 21 slices, TR/TE = 800/25 ms (MB factor = 3), and slice thickness = 6 mm. Inside the scanner, subjects were instructed to stay awake and keep their eyes closed, receiving verbal cues through headphones. The protocol included four fMRI sessions: a 15-minute conscious baseline, a 30-minute session during and after propofol infusion, and a final 15-minute recovery baseline. In the group analysis (e.g., Fig. 2C), first and last 5 minutes of unresponsive period was excluded from state-wise averaging to rule out transient effect.

### Dataset-2

The experimental protocol received approval from the IRB at the University of Michigan, ensuring all procedures adhered to applicable guidelines and regulations. The study enrolled thirty healthy individuals (10 males and 20 females, aged between 18 and 38 years, all right-handed). Each participant provided informed consent before participating and received compensation upon completion of the study. Three participants were excluded due to technical issues of MRI scanner or excessive movement during scanning. To protect participants’ privacy, stringent confidentiality measures were implemented. Each participant was allocated a unique code number at the initial point of contact, which was consistently used as the sole identifier for their specimen samples, as well as their behavioral, physiological, and MRI data, ensuring anonymity throughout the research process.

The criteria for including and excluding participants were defined as follows: Eligible participants were right-handed, healthy individuals classified as American Society of Anesthesiologists physical status 1, aged between 18 and 40, with a body mass index (BMI) of less than 30. Exclusion criteria encompassed any MRI contraindications such as potential pregnancy, severe obesity, the presence of metal in the body, claustrophobia, anxiety disorders, or any cardiopulmonary conditions; a history of neurological, cardiovascular, or respiratory diseases; significant head trauma involving loss of consciousness; any learning or developmental disabilities; a history of sleep apnea or pronounced snoring; any sensory or motor impairments that could affect study participation; gastroesophageal reflux disease; inability or unwillingness to refrain from alcohol for 24 hours before the MRI appointment; any history of illicit drug use or a positive result on a drug screening test; tattoos located on the head or neck area; known allergies to eggs; detectable intracranial abnormalities observed in T1-weighted MRI images; or any physical discomfort experienced during the fMRI scans.

The anesthetic administration and monitoring were consistent with that of Dataset-1, with the exception of certain specifics. The propofol infusion was manually adjusted to achieve stepwise target effect-site concentrations of 1.5, 2.0, 2.5, and 3.0 μg/mL. Each target concentration was sustained for a duration of 4 min, enabling us to incrementally titrate the anesthetic dosage to identify the threshold for loss of behavioral responsiveness (LOR). To reduce the potential for head motion-related artifacts, we kept the ESC at one level higher than the concentration associated with LOR for ≈ 32 min. For instance, if a participant exhibited LOR at a concentration of 2.0 μg/mL, the effect-site concentration was maintained at 2.5 μg/mL for the duration of the LOR phase. In rare cases (occurrence rate of ∼5.6%) where participants remained responsive at 3.0 μg/mL, the target concentration was escalated to 3.5 μg/mL and maintained at maximum of 4.0 μg/mL. Following this, the infusion was discontinued, allowing the propofol levels to diminish gradually. During this phase, participants were asked to engage in a behavioral task, rest, or listen to music as outlined below.

Throughout the experiment, a series of eight fMRI scans were performed, each lasting 16 minutes. These scans comprised a 16-minute resting-state (Rest1) and a 16-minute music-listening (Music1) session during the baseline period of wakefulness, followed by a 16-minute behavioral test conducted during the induction phase of propofol. After the participants had lost behavioral responsiveness, they underwent another set of 16-minute resting-state (Rest2) and music-listening (Music2) scans. This was succeeded by a 16-minute behavioral test during the emergence phase, which occurred post-termination of the propofol infusion. The final scans included a 16-minute resting-state (Rest3) and a 16-minute music-listening (Music3) session after the participants had regained behavioral responsiveness. Breaks ranging from 1 to 5 minutes were present between each scan. The entire imaging protocol and data collection process were completed within a total scanning session of 2.5 hours for each participant. In the group analysis (e.g., Fig. 2D), Rest1 and Music1 were defined as baseline state. Rest2 and Music2 were defined as LOR state. Rest3 and Music3 were defined as recovery state.

In the resting-state period, participants were instructed to lie still with their eyes closed, trying to remain awake while refraining from engaging in specific thoughts or movements. For the music-listening periods, they were asked to listen attentively to well-known tracks from four music genres—Jazz, Rock, Pop, and Country—with their eyes closed, remaining motionless and awake. The music tracks were edited to a consistent length of 4 minutes and played in a pseudo-random sequence. During the behavioral testing periods, participants were prompted to squeeze an MRI-compatible grip dynamometer (a rubber ball) at 10-second intervals for a total of 96 cycles. The cue to initiate each squeezing cycle was the spoken command “squeeze,” delivered through an MRI-compatible audiovisual system, with programming by E-Prime 3.0 software (Psychology Software Tools, Pittsburgh, PA). Headphone volume was set to ensure comfort during wakefulness and increased to 150% following the loss of responsiveness to ensure audibility. The participants’ grip strength was quantified in mmHg using the BIOPAC MP160 system coupled with AcqKnowledge software (V5.0). Analysis of the synchronization between the “squeeze” cues and the participants’ actual hand movements throughout the graded propofol administration allowed for the identification of the moments when subjects transitioned into and out of unresponsiveness. These transitions were marked by the first missed and successfully completed hand squeeze, respectively.

Data collection was conducted at the University of Michigan Hospital using a 3T Philips MRI scanner equipped with a standard 32-channel transmit/receive head coil. High-resolution anatomical images were captured using T1-weighted SPGR imaging techniques, configured with the following parameters: 170 sagittal slices, a slice thickness of 1.0 mm, a TR of 8.1 s, a TE of 3.7 ms, a flip angle of 8°, a FOV of 240 mm, and an image matrix size of 256 × 256. Functional brain imaging was performed employing a gradient-echo EPI sequence, detailed by the following specifications: 40 slices, TR/TE = 1400/30 ms by multiband acquisition (MB factor = 4), a slice thickness of 2.9 mm, an in-plane resolution of 2.75 × 2.75 mm, FOV of 220 mm, a flip angle of 76°, and an 80 × 80 image matrix.

### Dataset-3 (light and deep sedations)

The dataset has been previously published using analyses different from those applied here^58,59^. The IRB of Medical College of Wisconsin (MCW) approved the experimental protocol. The study involved fifteen healthy participants (nine males and six females, aged 19-35 years) who underwent sedation with propofol. The Observer’s Assessment of Alertness/Sedation (OAAS) scale was utilized to assess behavioral responsiveness levels. Participants were fully responsive to verbal stimuli with an OAAS score of 5 during the baseline and recovery conditions. In the light sedation condition, they exhibited lethargic responses to verbal commands, marked by an OAAS score of 4, and during deep sedation, they did not respond, with OAAS score ranging from 1 to 2. The target plasma concentrations for propofol varied among individuals (light sedation: 0.98 ± 0.18 μg/mL; deep sedation: 1.88 ± 0.24 μg/mL), reflecting the differences in personal sensitivity to the anesthetic. To maintain a steady state of sedation, the propofol plasma concentration was regulated by adjusting the infusion rate, balancing the drug’s accumulation and elimination, with manual adjustments informed by STANPUMP. Standard ASA-standard monitoring was conducted, which included tracking the electrocardiogram, blood pressure, pulse oximetry, and end-tidal CO_2_, along with prophylactic supplemental oxygen delivered through a nasal cannula. The study had to exclude one participant due to excessive movement, leaving 14 subjects available for subsequent analysis.

The acquisition of resting-state data encompassed four 15-minute scans, each corresponding to different conditions: baseline conscious, light sedation, deep sedation, and recovery. The images were captured using a 3T Signa GE 750 scanner (GE Healthcare, Waukesha, Wisconsin, USA) equipped with a standard 32-channel transmit/receive head coil. The scanner obtained whole-brain gradient-echo EPI sequences, detailed as follows: 41 slices per brain volume, TR = 2 s, TE = 25 ms, slice thickness = 3.5 mm, FOV = 224 mm, flip angle = 77°, and an image matrix size of 64 × 64. Alongside these functional scans, high-resolution anatomical images were captured for coregistration.

### Dataset-4 (light sedation and surgical level of propofol)

The dataset has been previously published using analyses different from those applied here^60^. The study received ethical clearance from the IRB of Huashan Hospital, Fudan University. The study enlisted twenty-six right-handed individuals (12 males and 14 females, aged between 27 and 64 years). Prior to their participation, informed consent was secured from all candidates, who subsequently received compensation post-experiment. These participants were all classified under the American Society of Anesthesiologists physical status I or II and were scheduled for elective surgery using a trans-sphenoidal approach to remove pituitary microadenomas. The diagnosis of these microadenomas was confirmed through radiological evaluations and plasma endocrine levels, identifying tumors less than 10 mm in diameter that did not extend beyond the sellar region. Exclusion criteria ensured that participants had no history of brain or significant organ dysfunction, or usage of neuropsychiatric medications, and were not at risk due to MRI contraindications like metal implants or vascular clips. However, three individuals were subsequently excluded from the analysis due to excessive movement, leaving data from 23 participants for further examination.

Before the study, participants were required to abstain from solid foods for a minimum of 8 hours and from liquids for 2 hours. Throughout the fMRI session, continuous monitoring was conducted for blood pressure, electrocardiography, pulse oximetry, and the partial pressure of CO_2_. An intravenous catheter was inserted into a vein on the right hand or forearm of the participants, through which propofol was administered to 17 out of 23 participants for light sedation and to all 23 participants for general anesthesia. The administration of propofol was regulated using a target-controlled infusion system, calibrated to maintain a steady effect-site concentration based on the Marsh model. For procedures under general anesthesia, remifentanil at a dose of 1.0 μg/kg and succinylcholine at 1.5 mg/kg were given to facilitate endotracheal intubation. The infusion was initiated at a concentration of 1.0 mg/mL, with incremental adjustments of 0.1 mg/mL until the desired effect-site concentration was achieved. A 5-minute interval was allowed for the propofol to evenly distribute across the compartments. The target propofol concentrations were maintained at 1.3 μg/mL for light sedation and 4.0 μg/mL for surgical level general anesthesia.

Behavioral responsiveness was evaluated using the Ramsay scale. Participants were categorized as fully conscious (Ramsay scores 1-2) when they provided a clear and prompt response to the command (“strongly squeeze my hand!”). If their response was clear but slow, they were considered to be mildly sedated (Ramsay scores 3-4). Those who did not respond were classified as deeply sedated or anesthetized (Ramsay scores 5–6). This specific verbal command of the Ramsay scale was administered twice during each evaluation phase. In conditions of conscious resting state and light sedation, participants were allowed to breathe spontaneously, receiving supplemental oxygen through a nasal cannula. Under general anesthesia, ventilation was assisted using intermittent positive pressure, ensuring a tidal volume between 8–10 mL/kg and a respiratory rate set between 10-12 breaths per minute. The entire procedure was supervised by two qualified anesthesiologists. To ensure auditory isolation and comfort during the fMRI scans, participants were equipped with earplugs and headphones.

Three 8-minute fMRI scans were conducted during conscious baseline, light sedation, and general anesthesia. To minimize head movement, participants’ heads were securely positioned within the scanner frame and supported by soft padding. They were instructed to lie back comfortably, remain supine, and keep their eyes closed, with an eye patch applied. During the resting-state scans, participants were advised to relax and not focus on any specific thoughts. The imaging was performed using a Siemens 3T MAGNETOM scanner equipped with a standard 8-channel head coil, capturing whole-brain gradient-echo EPI images under the following settings: 33 slices, a TR of 2000 ms, a TE of 30 ms, a 5 mm slice thickness, a 210 mm FOV, a 90° flip angle, and a 64 × 64 image matrix. High-resolution anatomical images were also obtained. For this study, the fMRI data collected during the conscious baseline and general anesthesia were particularly utilized.

### Dataset-5: Natural sleep

The dataset was acquired from the OpenNEURO database, shared by Pennsylvania State University with informed consent^36,37^. The original dataset included 33 healthy participants with wakefulness and three sleep stages (N1, N2, and N3). Sleep stage was identified based on EEG signatures determined by a Registered Polysomnographic Technologist^36,37^. MRI scans were conducted using a 3 Tesla Prisma Siemens Fit scanner, equipped with a Siemens 20-channel receive-array coil. The fMRI data was obtained through an EPI sequence, with a TR of 2.1 seconds, a TE of 25 ms, a slice thickness of 4 mm, comprising 35 slices, a FOV of 240 mm, and an in-plane resolution of 3 mm × 3 mm. One subject with head motion greater than 10 mm was excluded from the analysis. Only three out of 33 participants exhibited N3 sleep stages in their fMRI data, making analysis of this stage unreliable and leading to its exclusion. For ISD analysis, we excluded additional four subjects who had any of the stages (awake, N1, and N2) shorter than 1 minute (*n* = 28). For metastability and complexity analysis, we excluded 12 subjects who had any of the stages (awake, N1, and N2) shorter than 5 minutes (*n* = 20).

### Dataset-HCP

The dataset was acquired from the S1200 Release of the WU-Minn Human Connectome Project (HCP) database that has been fully described elsewhere^61^. Participants were healthy young adults from 22 to 37 years old. Participants who completed two sessions of resting-state fMRI scans (Rest1 and Rest2) were selected (*n* = 1009). Data collection utilized multiband EPI via a customized Siemens 3T MR scanner (Skyra system). Each scanning session included two sequences, each with different phase encoding directions (left-to-right and right-to-left), lasting 14 minutes and 33 seconds, with a TR of 720 ms, a TE of 33.1 ms, and a voxel size of 2-mm isotropic. The sequences from each session were merged, totaling 29 minutes and 6 seconds of data per session. This approach of merging data from opposite phase encodings aimed to neutralize any biases introduced by the direction of phase encoding. The denoised volumetric data, processed through ICA-FIX, were obtained from the online HCP database. Further details on the resting-state fMRI data collection and preprocessing steps are documented in other publications^62,63^. Static functional connectivity (FC) matrices of the Rest1 period produced by Pearson correlation were used for the analysis.

### fMRI data preprocessing and regional time course extraction

The preprocessing of fMRI data adhered to a comprehensive protocol established in our previous studies^48,56,64^. The AFNI software suite (linux_ubuntu_16_64; http://afni.nimh.nih.gov/) was utilized. Steps included: (1) Slice timing correction; (2) Rigid head motion correction/realignment; (3) Frame-wise scrubbing of head motion; (4) Coregistration with T1 anatomical images; (5) Spatial normalization into Talaraich stereotactic space resampled to 3 x 3 x 3 mm; (6) Band-pass filtering to 0.01 – 0.1 Hz while regressing out unwanted noise such as drifts or motion artifacts, and physiological noise from white matter and cerebrospinal fluid signals; (7) Spatial smoothing with 6 mm FWHM isotropic Gaussian kernel; (8) Temporal normalization to zero mean and unit variance. Cortex is defined by 400 cortical areas according to a well-established cortical parcellation scheme^65,66^. Subcortex is defined by 50 areas derived from a Subcortical Atlas^67^. The fMRI time courses were extracted from those 450 regions of interest (ROIs).

### Sliding window analysis

We adopted a sliding window approach for dynamic analysis. We chose a window size of 4 minutes with a 1-TR time step to reflect the dynamic changes in brain activity caused by the anesthetics instead of intrinsic variabilities^68^. However, we verified that the performance of ISD in discriminating awake and LOR states was stable across a range of window sizes from 0.5 to 8 minutes (Fig. 3D and Extended Data Fig. 4D). Pearson correlation was used to construct FC matrices.

### The use of global signal regression (GSR)

To assess different properties of functional brain networks, we utilized global signal regression (GSR). GSR is an effective way to mitigate confounding effects of motion that can affect the accuracy of network topology measurements. Moreover, since GSR artificially removes the global temporal coordination of fMRI signals, it can act as an effective way to regress the ‘integration’ of the network and reveal the underlying topological architecture (e.g., segregation), independent of global connectivity. Therefore, we utilized BOLD data that had not been subjected to GSR (noGSR) for calculating integration. For the computation of segregation, we utilized GSR-applied data.

### Multi-level efficiency and multi-level clustering coefficient

We propose novel measures, namely, multi-level efficiency and multi-level clustering coefficient as proxies for network integration and segregation, respectively. Refer to Extended Data Fig. 8 for an overview of the calculation procedures. An FC matrix was binarized at a given threshold *t* ranging from 0 to 1 (with an interval of 0.01). Then the global efficiency was calculated from the binary matrix:

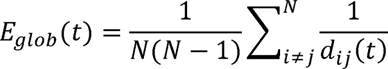

Where *d_ij_ (t)* is the shortest path length between nodes *i* and *j* when binarized at threshold *t*, and *N* is the total number of ROIs (= 450). Also, global clustering coefficient was calculated by averaging the local clustering coefficient:

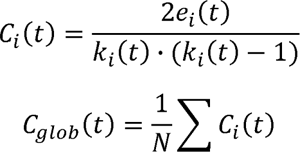

where *k_i_ (t)* and *e_i_ (t)* are the degree of node *i* and the number of edges between the neighbors of node *i* when binarized at threshold *t*.

Then, the multi-level efficiency (i.e. integration) and multi-level clustering coefficient (i.e. segregation) were determined by calculating the areas under the curve of the global efficiency and clustering coefficient values across the threshold range.

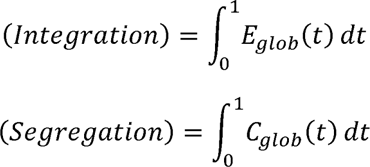

This method allows for a nuanced analysis of FC matrices by assessing network properties at various thresholds, thereby capturing the complexity of brain networks beyond single-scale approaches. Lastly, integration-segregation difference (ISD) was defined as the multi-level efficiency subtracted by multi-level clustering coefficient. This multi-thresholding approach is comparable with a previously suggested method of clustering coefficient calculation for complete weighted network^39^.

As both global efficiency and clustering coefficient are affected by density or connectivity of the FC matrix, we evaluated possible correlations of integration and segregation with connectivity values (i.e., mean value of positive edges) of FC matrices with and without GSR (Extended Data Fig. 9). High Spearman correlations were observed between integration and noGSR connectivity, and between segregation and GSR connectivity. This agrees with intuitive concept of integration and segregation, as integration can be understood as the amount of common global signal generated by a neural system. After GSR, the remaining signal likely reflects more localized, within-module connectivity^18^, as the global signal has been removed. Increased connectivity after GSR suggests enhanced strength of these local, segregated networks, indicating a shift towards greater segregation. Therefore, this demonstrates that our metrics effectively captured the integration and segregation of a given network.

### Brain network visualization

Brain plots were generated using BrainNet Viewer (https://www.nitrc.org/projects/bnv), in which connections stronger than *r* = 0.75 are shown. Topological profile of brain networks was visualized with the open-source software Gephi 0.10.1 (https://gephi.org/), in which the top 10% of connections are depicted using the ForceAtlas2 algorithm.

### Modularity and participation coefficient

Modularity is a measure that characterizes how well a network can be divided into modules, which is widely used metric for segregation. Modularity was defined by:

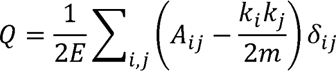

where *E* is total number of edges and *k_i_* is the degree of node *i*. The *δ_ij_* is 1 if *i* and *j* belong to same network, and otherwise 0. For module assignment, predefined 8 networks were used.

Participation coefficient quantifies between-module strength and has been widely used to measure network integration^15,18,25^. The participation coefficient of each node was calculated as:

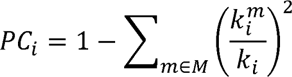

where *m* is the network in which *i* does not belong, and *k_i_^m^* is the sum of connections between node *i* and nodes in system *m*^18^. The average value of *PC_i_* across the nodes was then used as a proxy for network integration.

### Cartographic profiling

Cartographic profiling involves constructing 2D joint histograms of the participation coefficient and within-module z-score from every time frames and clustering these histogram profiles into two clusters, i.e., predominantly integrated and segregated states^25,69^. We followed the same procedure while using predefined 8-network assignment for participation coefficient and within-module z-score calculation. The k-means clustering (*k* = 2) was performed for each subject and was replicated 500 times with random initialization. The cluster with the higher average participation coefficient was assigned as the predominantly integrated state, while another cluster was labelled as the predominantly segregated state.

### System segregation

System segregation quantifies the strength of within-system connections in relation to between-system connections^18^. It is defined as

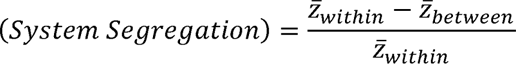

Where *Z̄_within_* is the mean value of intra-network Fisher z-transformed correlation values and *Z̄_between_* is the mean value of inter-network Fisher z-transformed correlation values. Following the original definition^18^, negative weights were excluded from calculation. Each node was assigned to a system based on the predefined 8 networks.

### Estimation of temporal regimes of ISD changes

To increase the statistical power, Datasets 1 and 2 were combined (total *n* = 45). For induction, the second scan from Dataset-1 and third scan from Dataset-2 were used. For emergence, the third scan from Dataset-1 and sixth scan from Dataset-2 were used. Each segment was time-locked to the onset of transition. If the lengths of the data before or after transition were shorter than 8 min, the empty values were filled with the mean ISD trends to ensure continuity of smoothing. Then each curve was spline-smoothed with the smoothing parameter of 10^-6^, which reasonably preserves general trend while suppressing transient fluctuation. The slope was estimated by getting time derivative of the smoothed curves. For each second, the distribution of 45 slope values was tested against zero median (Wilcoxon signed-rank test, FDR-corrected *p* < 0.05). Log_10_ of *p*-values were applied to moving average of a 4-minute window.

The goodness of fit was quantified as normalized mean absolute error (NMAE), defined as:

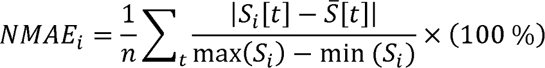

Where *S_i_*[*t*] is spline-smoothed ISD curve of *i*th subject and *S*[*t*] is the group average of spline smoothed curves.

### Training and evaluation of machine learning models

Integration and segregation values were extracted from the dynamic FC matrices of nine networks (whole-brain and eight pre-defined networks). For model comparison, we aggregated the integration and segregation across all networks, culminating in a feature set of 18 measurements. For training, data points from stable awake and LOR states (Dataset 1: Baseline, Deep LOR (LOR excluding first and last 5 min), and Recovery; Dataset 2: Rest1, Music1, Music3, and Rest3) were used. Data points close to state transitions were not seen to the model during training. Considering the class imbalance in the training set, non-overlapping 20000 data points were randomly selected from each class (awake and LOR) during training for all five models (total 40000 training data points).

The performance of models was evaluated by the AUC of the ROC curve and balanced accuracy (the mean of sensitivity and specificity). We adopted a leave-one-subject-out (LOSO) strategy for cross-validation and hyperparameter selection^70^. This method involved sequentially excluding one subject from the training process and utilizing the omitted subject’s data as the validation set. To ensure the generalizability of the model, hyperparameters chosen from Dataset-1 were also applied to Dataset-2.

For random forest (RF), the number of trees is set at 100. Trees are allowed to grow until all leaves are pure. For each split, five features are randomly sampled, equivalent to the ceiling of square root of the number of total features. For support vector machine (SVM), we selected a radial basis function (RBF) kernel. The regularization parameter *C* was set to 1.0. Kernel scale parameter was heuristically chosen by ‘fitcsvm’ function in MATLAB. For artificial neural network (ANN), we implemented a pattern recognition network with two hidden layers (10 and 2 neurons each), utilizing a scaled conjugate gradient training function and cross-entropy as the performance function. The learning rate and momentum was set as 0.01 and 0.9, respectively. We employed L2 regularization to prevent overfitting. Our ANN was configured to use rectified liner units (ReLUs) for activation and included dropout and early stopping to enhance performance. For *k*-nearest neighbor (KNN) model, we selected the optimal value of k = 5 via cross-validation. We employed the Euclidean distance metric due to its effectiveness in continuous data spaces. For logistic regression, we employ a logit link function to predict consciousness states.

### Metastability

We calculated the moving standard deviation of Kuramoto order parameters as a proxy to quantify metastability in the neural system^28^. Since GSR removes the entire global synchronization, the noGSR data was used for metastability calculation. The BOLD time series of ROIs were transformed to coupled oscillators via Hilbert transform. The Kuramoto order parameter (*R*) was defined as:

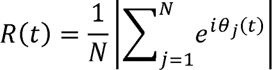

Where *θ_j_* is the instantaneous phase of *j*th ROI. Metastability was quantified as the moving standard of the Kuramoto order parameter (*σ_R_*) with the sliding window of 4 min.

### Pattern complexity

We introduce a novel metric to quantify the complexity of the BOLD signal pattern. The parcellated BOLD signal can be considered as 450-dimensional vectors, which can be clustered by *k*-means clustering method, a commonly used unsupervised machine learning algorithm. The clustering was performed at subject-level, in which the value of *k* ranged from 3 to 100. To ensure convergence, the clustering was replicated 100 times for each *k*. Considering the data’s high dimensionality, correlation distance was used as a distance metric. This converts the BOLD signal into pattern sequence.

Subsequently, we calculated three different aspects of complexity with the sliding window of 4 minutes. First, the entropy of pattern occurrence was calculated as:

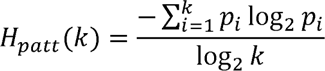

where *p_i_* is the occurrence rate of *i*th pattern within the given sequence. This quantifies how evenly every pattern appears. Second, we transformed the pattern sequence into *k*-by-*k* transition matrix and calculated its entropy as:

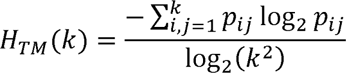

where *p_ij_* is the occurrence rate of *i*-to-*j* transition. Third, we used an effort-to-compress (*ETC*) to quantify the temporal complexity of the pattern sequence^33,35^. *ETC* utilizes a lossless compression through non-sequential recursive pair substitution and describes how many substitutions are needed to fully describe the categorical sequence data. Following the consideration of the previous study^33^, we normalized the raw *ETC* by dividing with, where *L* is the length of the series. We combined them by averaging their normalized AUC. We named this composite measure as *pattern complexity* (*C_patt_*).

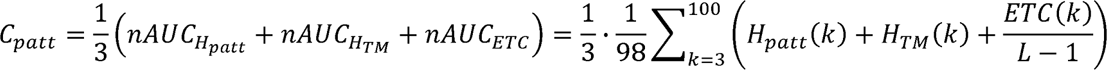

The strong mutual correlations among the entropy of pattern occurrence (*H_patt_*), transition matrix (*H_TM_*), and the effort-to-compress (*ETC*) (Extended Data Fig. 7) suggest that our pattern complexity measure (*C_patt_*) is a robust indicator of the brain’s dynamical state. We confirmed that GSR had negligible effect on the pattern complexity.

### Statistics and reproducibility

For all statistical analyses in Fig. 2D,E,F, 6F, 7E, 8A,B,D, measurements during three conditions (baseline, LOR, and recovery for anesthesia data; awake, N1, and N2 sleep for sleep data) were statistically compared using Friedman’s ANOVA and post-hoc Wilcoxon test (two-sided). The statistical significance was set at FDR-corrected *p* < 0.05. All statistical tests used in this study are two-sided. In Fig. 4, 45 slope values for each time point were tested against zero median by Wilcoxon signed-rank test, FDR-corrected *p* < 0.05. Multiple comparison was conducted for the entire networks and all time points. In Fig. 6I, 7H, we employed a dominance analysis^31^ to investigate the relationship between metastability/complexity (the response variables) and two key predictors: integration and segregation.

While individual fMRI data underwent standard motion correction (e.g., motion correction, scrubbing, regressing out motion profiles), we conducted additional group-level control analyses to account for subject- or state-specific motion effects on ISD, metastability, and pattern complexity. Linear regression, using state-average FD values as a second level regressor, isolated fMRI-derived measures from possible motion effects. ANOVA and post-hoc tests on these adjusted values were performed to examine the robustness of our findings (Table S1-3).

The key finding of the paper, i.e., decrease of ISD under LOR induced by propofol anesthetics or natural sleep, was first found in Dataset-1, which was reproduced independently in Datasets 2 to 5. Another key finding of the paper, near-zero value of ISD of healthy awake brain was first found in Dataset-1 and replicated independently in Datasets 2 to 5 and Dataset-HCP. Relationships among metastability, complexity, integration, and segregation were first found in Dataset-1 and independently replicated in Dataset-2 and Dataset-5.

## Data Availability

Data used in the analyses will become publicly available from Zenodo repository. Source data to plot the figures will be provided with this paper. The natural sleep fMRI dataset is available from OpenNEURO (https://openneuro.org/datasets/ds003768/versions/1.0.11). The HCP dataset is available from online repository (https://www.humanconnectome.org/).

## Code Availability

Custom-built code for ISD, metastability, and pattern complexity calculations is available at Github (https://github.com/janghw4/multilevel-integration-segregation-balance).

## Acknowledgments

This work was supported by the National Institute of General Medical Sciences of the National Institutes of Health grants 2R01-GM103894 (to A.G.H. and Z.H.). The content is solely the responsibility of the author and does not necessarily represent the official views of the National Institutes of Health.

## Author Contributions Statement

H.J. and Z.H. conceptualized this work. H.J. developed the analytical methods, analyzed the data, prepared the figures, and drafted the manuscript. Z.H. collected and co-analyzed the data. Z.H., G.A.M. and A.G.H. interpreted the data and edited the manuscript.

---

**Extended Data Figure 1.**
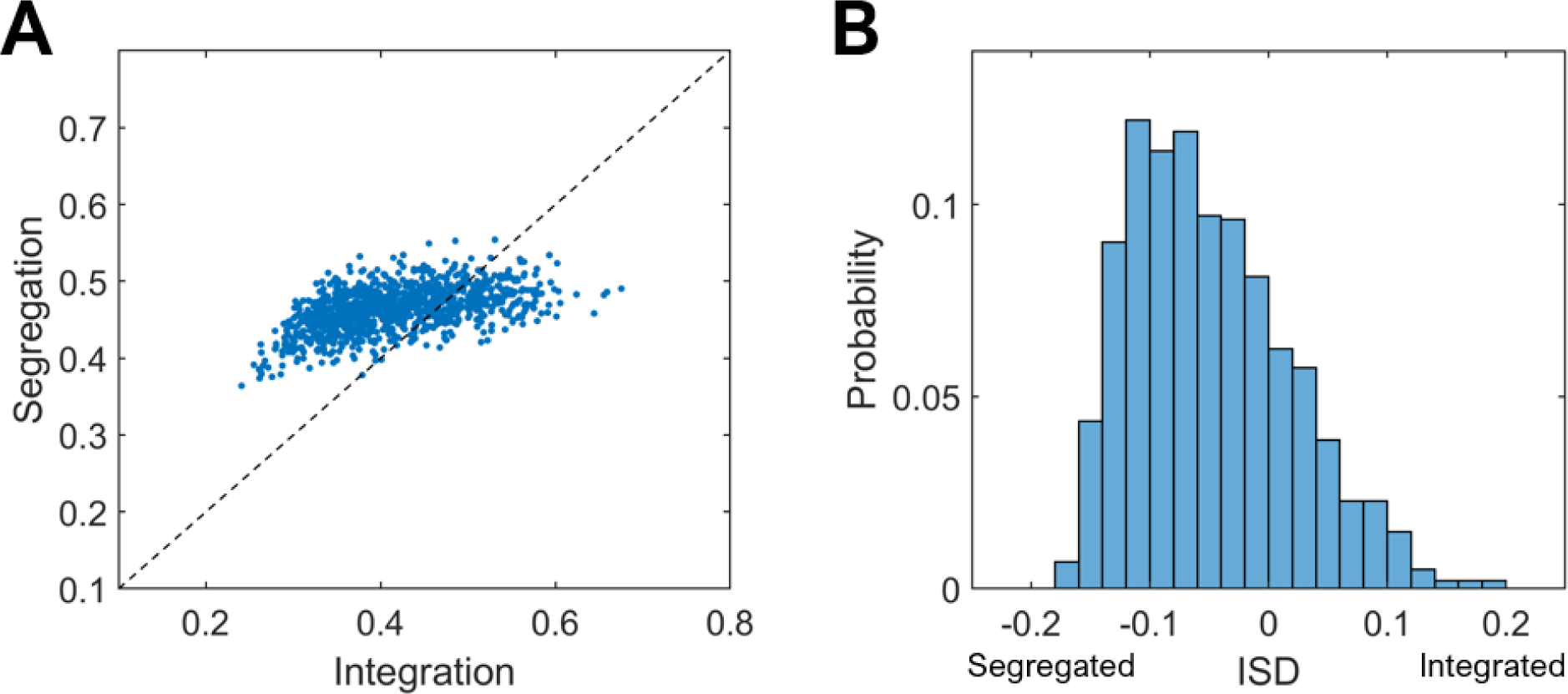
(A) Scatter plot of integration and segregation and (B) histogram of ISD values for individuals in Dataset-HCP (*n* = 1009). Dashed line indicates *y* = *x*. The values of integration and segregation were 0.41 ± 0.08 (mean ± SD) and 0.46 ± 0.03, respectively. The difference between integration and segregation (ISD) was –0.05 ± 0.07.

**Extended Data Figure 2.**
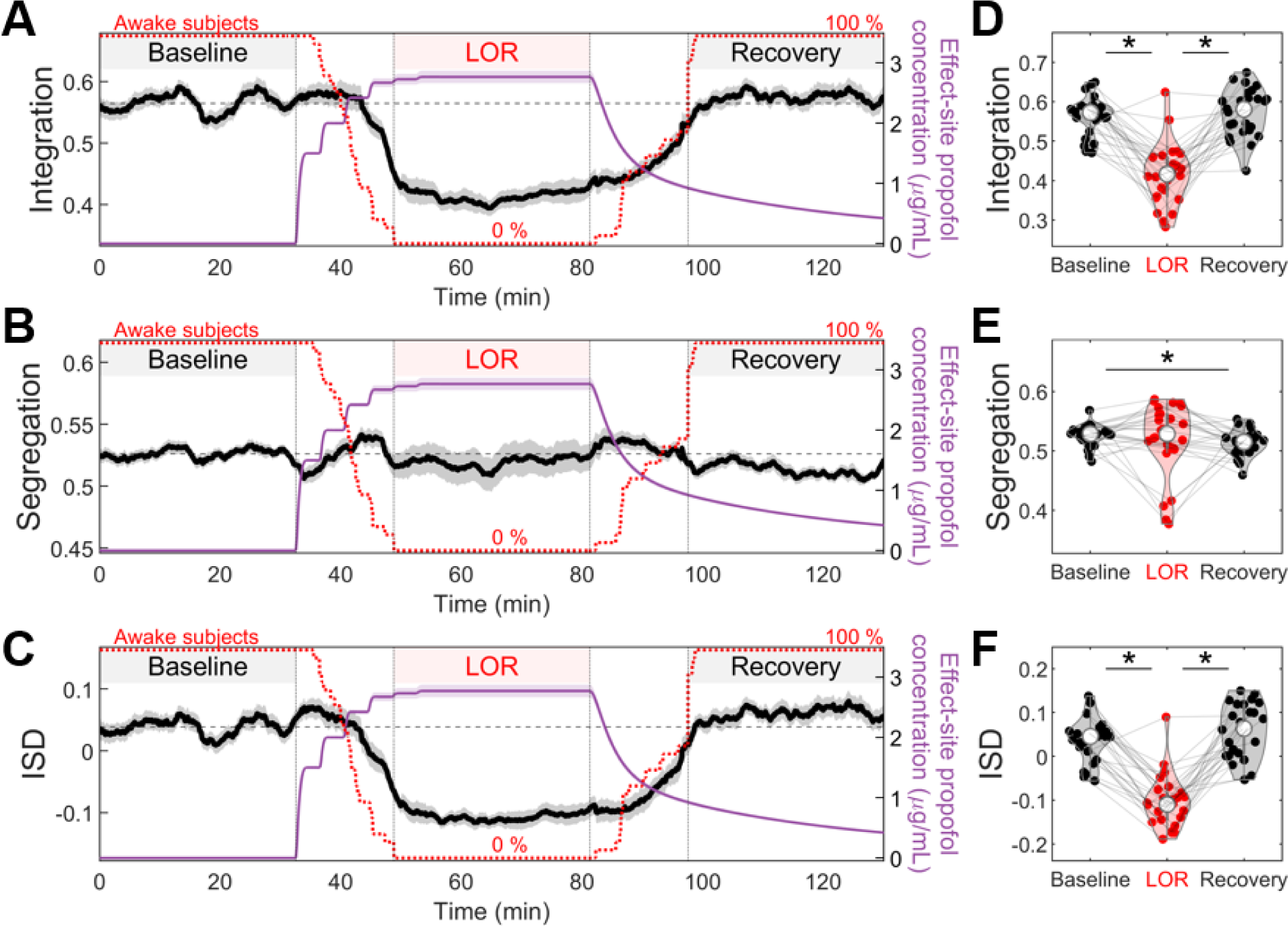
(A-C) Average time course of integration, segregation, and ISD for Dataset-2 (*n* = 26). Horizontal dashed lines are mean values during baseline. Shaded areas represent ± SEM. Sliding window size was 4 minutes. (D-F) Average values for integration, segregation, and ISD during baseline, loss of responsiveness (LOR), and recovery. Each point represents an individual subject, linked by grey lines. Median values are marked by white circles. Asterisks denote statistically significant differences (paired two-sided Wilcoxon signed rank test) with FDR-corrected p < 0.05. ISD showed significant changes across states (Friedman’s ANOVA: *p* = 6.79 × 10^-9^, Post-hoc Wilcoxon test: baseline vs. LOR *p* = 1.77 × 10^-5^, LOR vs. recovery *p* = 1.77 × 10^-5^, baseline vs. recovery *p* = 0.1014).

**Extended Data Figure 3.**
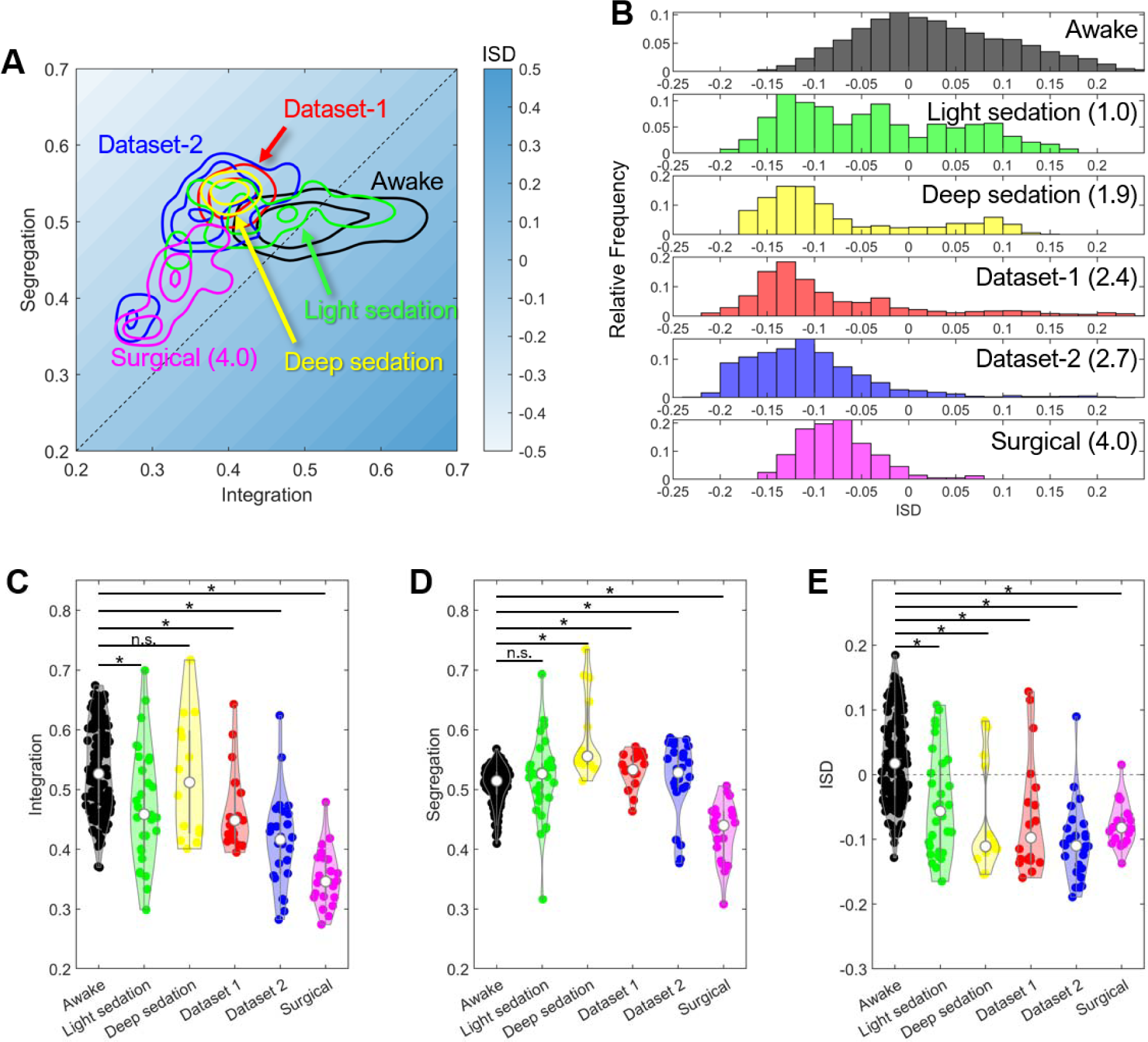
(A) Contour plots of various anesthetic conditions in integration-segregation parameter space. (B) Histograms of ISD for various anesthetic dosages. Numbers in parenthesis correspond to approximate effect site concentration (ESC) of propofol. Violin plots of (C) integration, (D) segregation, and (E) ISD averaged for each subject. Asterisks denote statistically significant differences (two-sided Mann-Whitney U-test) with FDR-corrected *p* < 0.05.

**Extended Data Figure 4.**
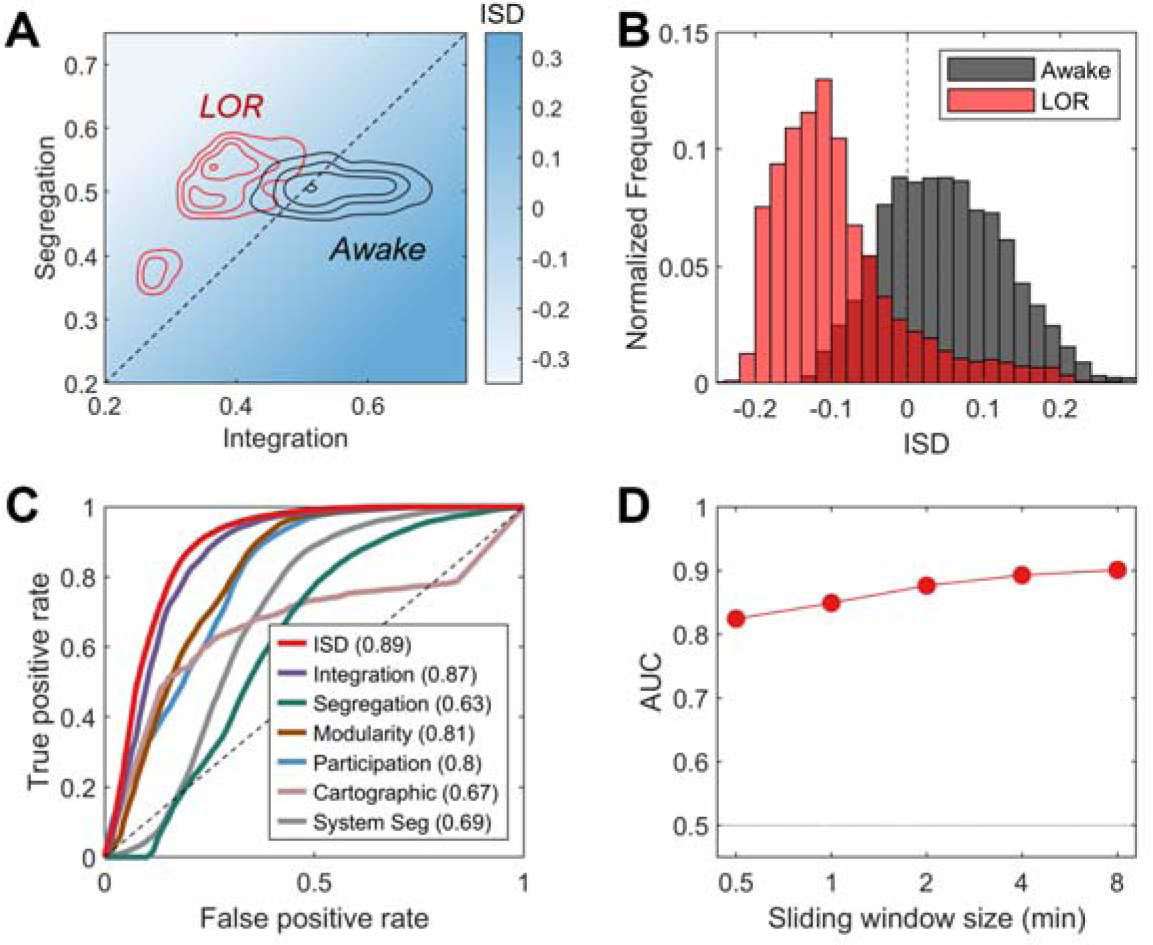
Replication of performance analysis with Dataset-2. (A) A contour density plot illustrating the distribution of awake (gray) and LOR (red) states across integration and segregation parameters. (B) Histograms of ISD value distribution for awake (gray) and LOR (red) conditions. (C) ROC curve analysis comparing the performance of ISD and other network metrics, quantified by AUC of ROC curves. Metrics include ISD, integration, segregation, modularity, participation coefficient, cartographic method, and system segregation. (D) The relationship between the performance of ISD (as measured by AUC) and the size of sliding window.

**Extended Data Figure 5.**
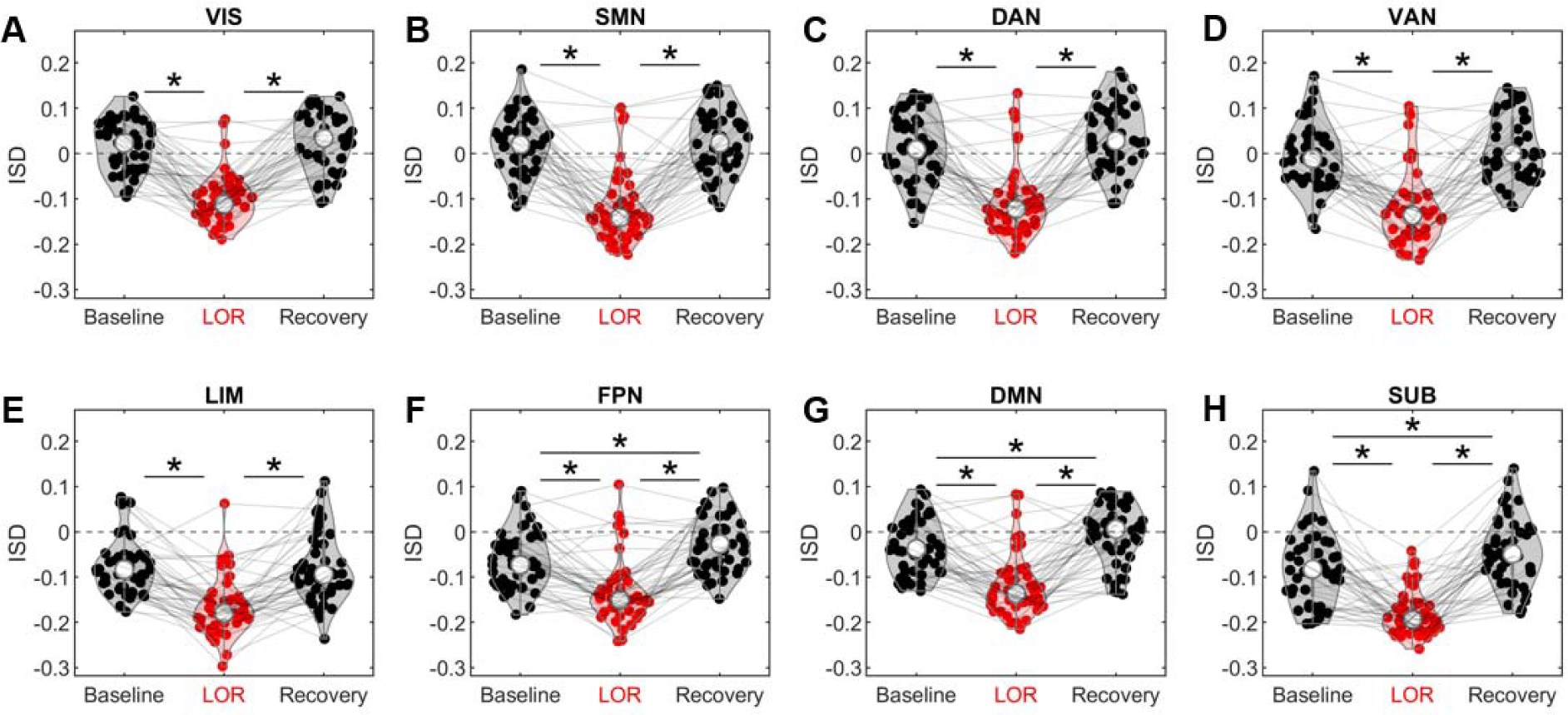
Network-level ISD during baseline, loss of responsiveness (LOR), and recovery. Each point represents an individual subject, linked by grey lines for clarity. Median values are marked by white circles. Asterisks denote statistically significant differences (paired two-sided Wilcoxon signed rank test) with FDR-corrected *p* < 0.05. Datasets 1 and 2 were merged (*n* = 45).

**Extended Data Figure 6.**
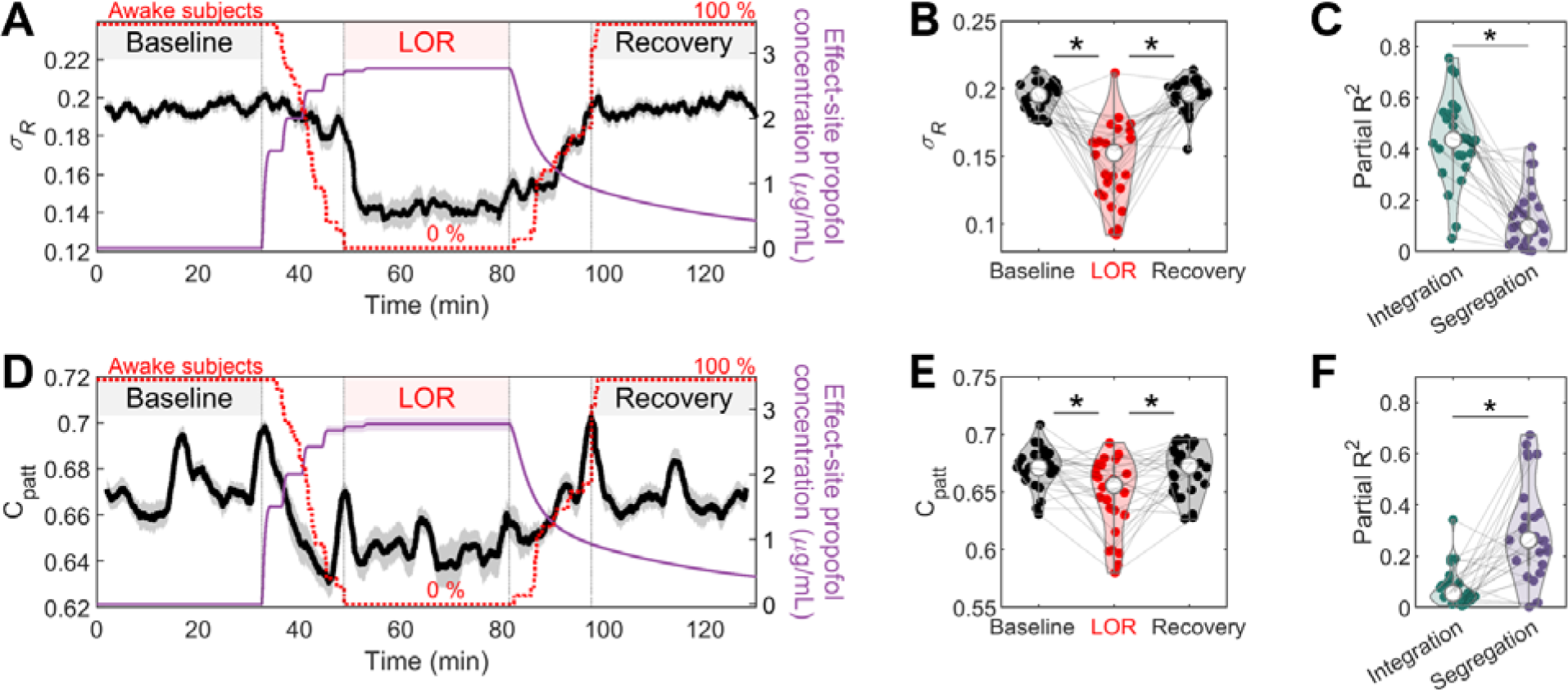
(A) A time course of metastability during loss and recovery of responsiveness in Dataset-2. (B) State-averaged values of metastability. Metastability showed significant changes across states (Friedman’s ANOVA: *p* = 1.47 × 10^-7^, Post-hoc Wilcoxon test: baseline vs. LOR *p* = 2.50 × 10^-5^, LOR vs. recovery *p* = 2.50 × 10^-5^, baseline vs. recovery *p* = 0.3809). (C) Partial *R*^2^ from a dominance analysis relating metastability to integration and segregation. Partial *R*^2^ values for integration and segregation were 0.4355 ± 0.1673 and 0.1228 ± 0.1144 respectively (*p* = 9.34 x 10^-6^). (D) A time course of pattern complexity in Dataset-2. (E) State-averaged values of pattern complexity. Pattern complexity showed significant changes across states (Friedman’s ANOVA: *p* = 0.0151, Post-hoc Wilcoxon test: baseline vs. LOR *p* = 0.0036, LOR vs. recovery *p* = 0.0102, baseline vs. recovery *p* = 0.3673). (F) Partial *R*^2^ from a dominance analysis relating complexity to integration and segregation. Partial *R*^2^ values for integration and segregation were 0.0769 ± 0.0753 and 0.2950 ± 0.1940 respectively (*p* = 1.19 x 10^-4^). The asterisks denote statistically significant differences (paired two-sided Wilcoxon signed rank test) with *p* < 0.05.

**Extended Data Figure 7.**
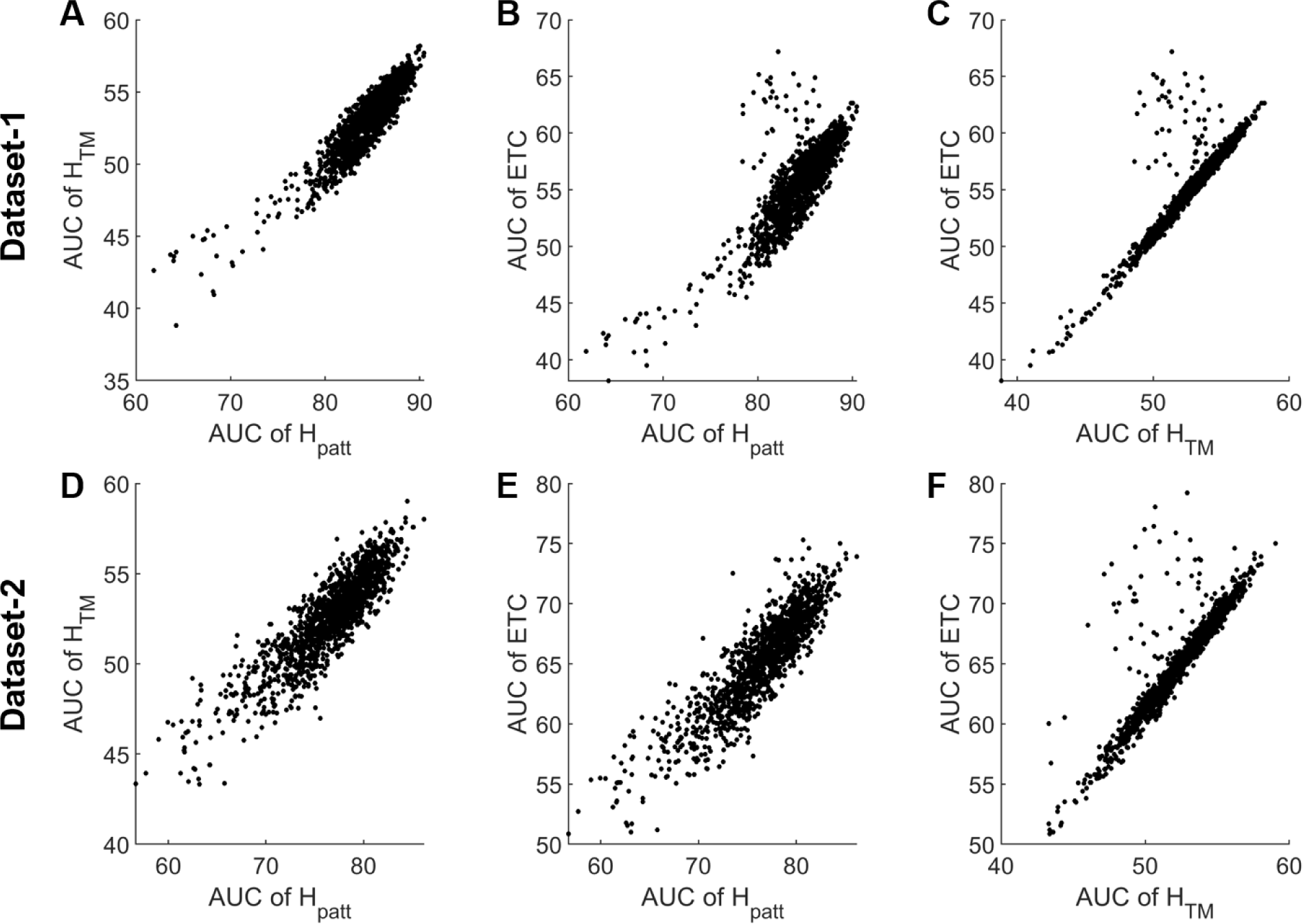
(A-C) Scatter plots of areas under the curve (AUC) of (A) *H_patt_* vs. *H_TM_*, (B) *H_patt_* vs. *H_TM_*, (C) *H_TM_* vs. *ETC* for Dataset-1. (D-F) Identical analysis for Dataset-2.

**Extended Data Figure 8.**
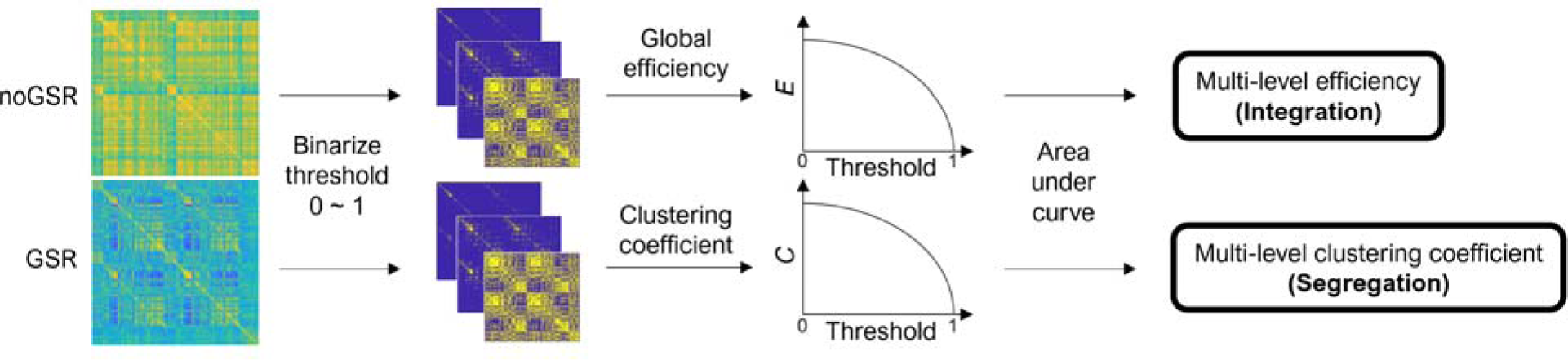
Overview of a procedure to calculate multi-level efficiency (integration) and multi-level clustering coefficient (segregation).

**Extended Data Figure 9.**
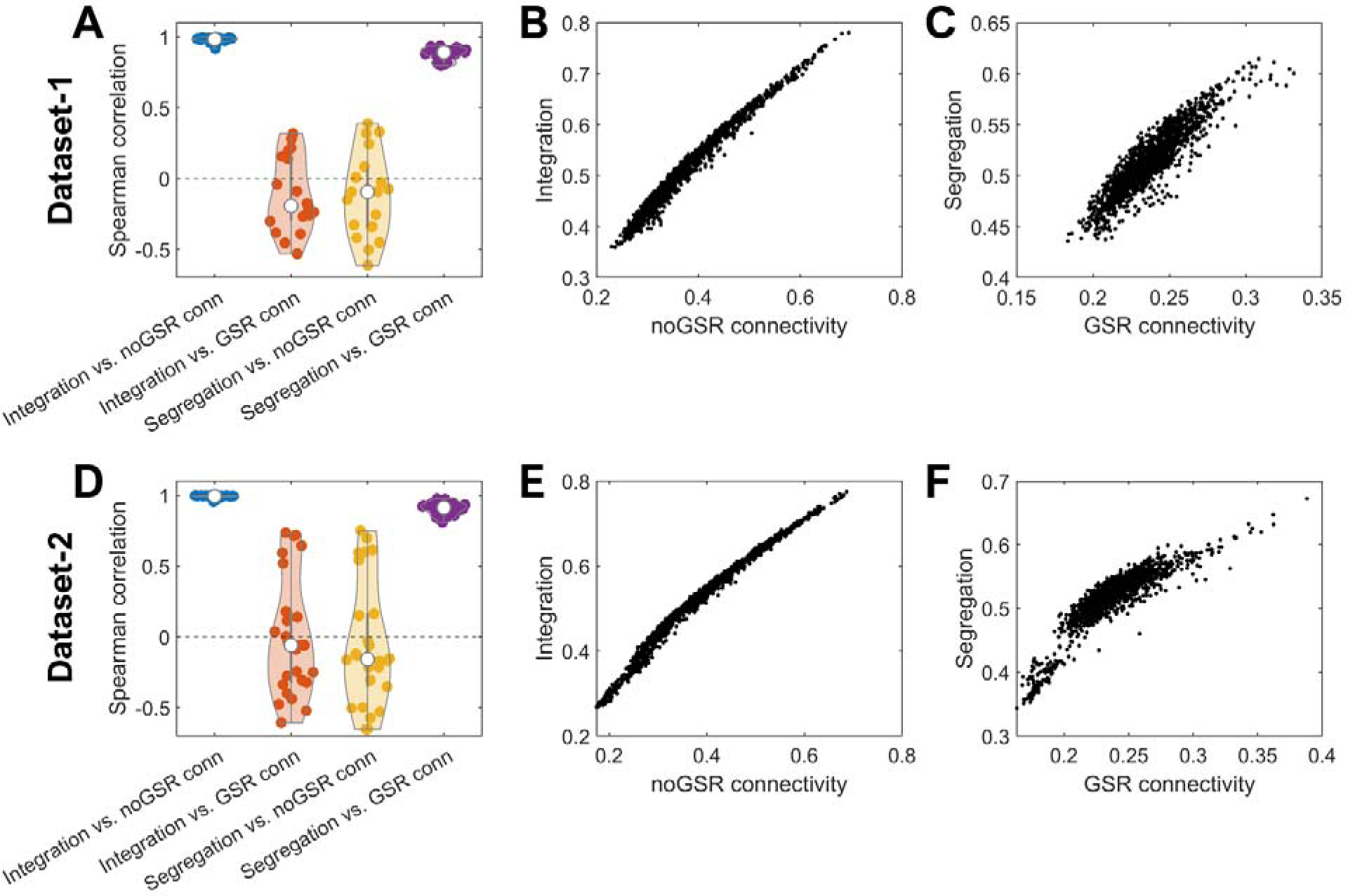
(A) Spearman correlation of integration and segregation with mean connectivity (average of positive edge values) of noGSR and GSR functional connectivity (FC) matrices for Dataset-1. (B-C) Scatter plots of (B) integration vs. noGSR connectivity and (C) segregation vs. GSR connectivity. (D-F) Identical analysis for Dataset-2.

