## Supplementary material for "Measuring the dynamic balance of integration and segregation underlying consciousness, anesthesia, and sleep": Table S

**Table S1.** Additional statistics for ISD analysis in Dataset-1 and Dataset-2 (see Fig. 2 and Extended Data Fig. 2).

| **Before additional  regression of  state-averaged FD** | **Dataset-1, Friedman’s ANOVA** | | | | | |
| --- | --- | --- | --- | --- | --- | --- |
|  | Source | SS | df | MS | Chi-sq | Prob>Chi-sq |
|  | Columns | 12.3158 | 2 | 6.15789 | 12.32 | 0.0021 |
|  | Error | 25.6842 | 36 | 0.71345 |  |  |
|  | Total | 38 | 56 |  |  |  |
|  | **Dataset-1, Post-hoc Wilcoxon test** | | | | | |
|  | Condition | | Raw p-value | | FDR-corrected p-value | |
|  | Baseline vs. LOR | | 0.0033 | | 0.0050 | |
|  | Recovery vs. LOR | | 0.0004 | | 0.0012 | |
|  | Baseline vs. Recovery | | 0.1165 | | 0.1165 | |
|  | **Dataset-2, Friedman’s ANOVA** | | | | | |
|  | Source | SS | df | MS | Chi-sq | Prob>Chi-sq |
|  | Columns | 37.6154 | 2 | 18.8077 | 37.62 | 6.79 x 10^-9^ |
|  | Error | 14.3846 | 50 | 0.2877 |  |  |
|  | Total | 52 | 77 |  |  |  |
|  | **Dataset-2, Post-hoc Wilcoxon test** | | | | | |
|  | Condition | | Raw p-value | | FDR-corrected p-value | |
|  | Baseline vs. LOR | | 1.18 x 10^-5^ | | 1.77 x 10^-5^ | |
|  | Recovery vs. LOR | | 8.30 x 10^-6^ | | 1.77 x 10^-5^ | |
|  | Baseline vs. Recovery | | 0.1014 | | 0.1014 | |
| **After additional  regression of  state-averaged FD** | **Dataset-1, Friedman’s ANOVA** | | | | | |
|  | Source | SS | df | MS | Chi-sq | Prob>Chi-sq |
|  | Columns | 7.6842 | 2 | 3.84211 | 7.68 | 0.0214 |
|  | Error | 30.3158 | 36 | 0.84211 |  |  |
|  | Total | 38 | 56 |  |  |  |
|  | **Dataset-1, Post-hoc Wilcoxon test** | | | | | |
|  | Condition | | Raw p-value | | FDR-corrected p-value | |
|  | Baseline vs. LOR | | 0.0112 | | 0.0169 | |
|  | Recovery vs. LOR | | 0.0015 | | 0.0044 | |
|  | Baseline vs. Recovery | | 0.0910 | | 0.0910 | |
|  | **Dataset-2, Friedman’s ANOVA** | | | | | |
|  | Source | SS | df | MS | Chi-sq | Prob>Chi-sq |
|  | Columns | 29.1538 | 2 | 14.5769 | 29.15 | 4.67 x 10^-7^ |
|  | Error | 22.8462 | 50 | 0.4569 |  |  |
|  | Total | 52 | 77 |  |  |  |
|  | **Dataset-2, Post-hoc Wilcoxon test** | | | | | |
|  | Condition | | Raw p-value | | FDR-corrected p-value | |
|  | Baseline vs. LOR | | 2.10 x 10^-5^ | | 3.15 x 10^-5^ | |
|  | Recovery vs. LOR | | 1.18 x 10^-5^ | | 1.35 x 10^-5^ | |
|  | Baseline vs. Recovery | | 0.0775 | | 0.0775 | |

**Table S2.** Additional statistics for metastability analysis in Dataset-1 and Dataset-2 (see Fig. 6 and Extended Data Fig. 6A-C).

| **Before additional  regression of  state-averaged FD** | **Dataset-1, Friedman’s ANOVA** | | | | | |
| --- | --- | --- | --- | --- | --- | --- |
|  | Source | SS | df | MS | Chi-sq | Prob>Chi-sq |
|  | Columns | 15.4737 | 2 | 7.73684 | 15.47 | 0.0004 |
|  | Error | 22.5263 | 36 | 0.62573 |  |  |
|  | Total | 38 | 56 |  |  |  |
|  | **Dataset-1, Post-hoc Wilcoxon test** | | | | | |
|  | Condition | | Raw p-value | | FDR-corrected p-value | |
|  | Baseline vs. LOR | | 0.0089 | | 0.0134 | |
|  | Recovery vs. LOR | | 0.0005 | | 0.0016 | |
|  | Baseline vs. Recovery | | 0.1075 | | 0.1075 | |
|  | **Dataset-2, Friedman’s ANOVA** | | | | | |
|  | Source | SS | df | MS | Chi-sq | Prob>Chi-sq |
|  | Columns | 31.4615 | 2 | 15.7308 | 31.46 | 1.47 x 10^-7^ |
|  | Error | 20.5385 | 50 | 0.4108 |  |  |
|  | Total | 52 | 77 |  |  |  |
|  | **Dataset-2, Post-hoc Wilcoxon test** | | | | | |
|  | Condition | | Raw p-value | | FDR-corrected p-value | |
|  | Baseline vs. LOR | | 1.67 x 10^-5^ | | 2.50 x 10^-5^ | |
|  | Recovery vs. LOR | | 1.49 x 10^-5^ | | 2.50 x 10^-5^ | |
|  | Baseline vs. Recovery | | 0.3809 | | 0.3809 | |
| **After additional  regression of  state-averaged FD** | **Dataset-1, Friedman’s ANOVA** | | | | | |
|  | Source | SS | df | MS | Chi-sq | Prob>Chi-sq |
|  | Columns | 14 | 2 | 7 | 14 | 0.0009 |
|  | Error | 24 | 36 | 0.66667 |  |  |
|  | Total | 38 | 56 |  |  |  |
|  | **Dataset-1, Post-hoc Wilcoxon test** | | | | | |
|  | Condition | | Raw p-value | | FDR-corrected p-value | |
|  | Baseline vs. LOR | | 0.0269 | | 0.0403 | |
|  | Recovery vs. LOR | | 0.0043 | | 0.0128 | |
|  | Baseline vs. Recovery | | 0.1075 | | 0.1075 | |
|  | **Dataset-2, Friedman’s ANOVA** | | | | | |
|  | Source | SS | df | MS | Chi-sq | Prob>Chi-sq |
|  | Columns | 27 | 2 | 13.5 | 27 | 1.37 x 10^-6^ |
|  | Error | 25 | 50 | 0.5 |  |  |
|  | Total | 52 | 77 |  |  |  |
|  | **Dataset-2, Post-hoc Wilcoxon test** | | | | | |
|  | Condition | | Raw p-value | | FDR-corrected p-value | |
|  | Baseline vs. LOR | | 3.67 x 10^-5^ | | 5.51 x 10^-5^ | |
|  | Recovery vs. LOR | | 1.87 x 10^-5^ | | 5.51 x 10^-5^ | |
|  | Baseline vs. Recovery | | 0.3539 | | 0.3539 | |

**Table S3.** Additional statistics for pattern complexity analysis in Dataset-1 and Dataset-2 (see Fig. 7 and Extended Data Fig. 6D-F).

| **Before additional  regression of  state-averaged FD** | **Dataset-1, Friedman’s ANOVA** | | | | | |
| --- | --- | --- | --- | --- | --- | --- |
|  | Source | SS | df | MS | Chi-sq | Prob>Chi-sq |
|  | Columns | 28.7368 | 2 | 14.3684 | 28.74 | 5.75 x 10^-7^ |
|  | Error | 9.2632 | 36 | 0.2573 |  |  |
|  | Total | 38 | 56 |  |  |  |
|  | **Dataset-1, Post-hoc Wilcoxon test** | | | | | |
|  | Condition | | Raw p-value | | FDR-corrected p-value | |
|  | Baseline vs. LOR | | 0.0001 | | 0.0002 | |
|  | Recovery vs. LOR | | 0.0001 | | 0.0002 | |
|  | Baseline vs. Recovery | | 0.8092 | | 0.8092 | |
|  | **Dataset-2, Friedman’s ANOVA** | | | | | |
|  | Source | SS | df | MS | Chi-sq | Prob>Chi-sq |
|  | Columns | 8.3846 | 2 | 4.19231 | 8.38 | 0.0151 |
|  | Error | 43.6154 | 50 | 0.87231 |  |  |
|  | Total | 52 | 77 |  |  |  |
|  | **Dataset-2, Post-hoc Wilcoxon test** | | | | | |
|  | Condition | | Raw p-value | | FDR-corrected p-value | |
|  | Baseline vs. LOR | | 0.0012 | | 0.0036 | |
|  | Recovery vs. LOR | | 0.0068 | | 0.0102 | |
|  | Baseline vs. Recovery | | 0.3673 | | 0.3673 | |
| **After additional  regression of  state-averaged FD** | **Dataset-1, Friedman’s ANOVA** | | | | | |
|  | Source | SS | df | MS | Chi-sq | Prob>Chi-sq |
|  | Columns | 17.7895 | 2 | 8.89474 | 17.79 | 0.0001 |
|  | Error | 20.2105 | 36 | 0.5614 |  |  |
|  | Total | 38 | 56 |  |  |  |
|  | **Dataset-1, Post-hoc Wilcoxon test** | | | | | |
|  | Condition | | Raw p-value | | FDR-corrected p-value | |
|  | Baseline vs. LOR | | 0.0015 | | 0.0022 | |
|  | Recovery vs. LOR | | 0.0011 | | 0.0022 | |
|  | Baseline vs. Recovery | | 1 | | 1 | |
|  | **Dataset-2, Friedman’s ANOVA** | | | | | |
|  | Source | SS | df | MS | Chi-sq | Prob>Chi-sq |
|  | Columns | 5.6154 | 2 | 2.80768 | 5.62 | 0.0603 |
|  | Error | 46.3846 | 50 |  |  |  |
|  | Total | 52 | 77 |  |  |  |
|  | **Dataset-2, Post-hoc Wilcoxon test** | | | | | |
|  | Condition | | Raw p-value | | FDR-corrected p-value | |
|  | Baseline vs. LOR | | 0.0409 | | 0.0614 | |
|  | Recovery vs. LOR | | 0.0339 | | 0.0614 | |
|  | Baseline vs. Recovery | | 0.5506 | | 0.5506 | |

**Table S4.** Additional statistics for Dataset-5 analysis (see Fig. 8).

| **Integration** | **Friedman’s ANOVA** | | | | | |
| --- | --- | --- | --- | --- | --- | --- |
|  | Source | SS | df | MS | Chi-sq | Prob>Chi-sq |
|  | Columns | 10.2857 | 2 | 5.14286 | 10.29 | 0.0058 |
|  | Error | 45.7143 | 54 | 0.84656 |  |  |
|  | Total | 56 | 83 |  |  |  |
|  | **Post-hoc Wilcoxon test** | | | | | |
|  | Condition | | Raw p-value | FDR-corrected p-value | | |
|  | Awake vs. N2 | | 0.0215 | 0.0322 | | |
|  | N1 vs. N2 | | 0.0139 | 0.0322 | | |
|  | Awake vs. N1 | | 0.1329 | 0.1329 | | |
| **Segregation** | **Friedman’s ANOVA** | | | | | |
|  | Source | SS | df | MS | Chi-sq | Prob>Chi-sq |
|  | Columns | 2.7857 | 2 | 1.39286 | 2.79 | 0.2484 |
|  | Error | 53.2143 | 54 | 0.98545 |  |  |
|  | Total | 56 | 83 |  |  |  |
| **ISD** | **Friedman’s ANOVA** | | | | | |
|  | Source | SS | df | MS | Chi-sq | Prob>Chi-sq |
|  | Columns | 9.5 | 2 | 4.75 | 9.5 | 0.0087 |
|  | Error | 46.5 | 54 | 0.86111 |  |  |
|  | Total | 56 | 83 |  |  |  |
|  | **Post-hoc Wilcoxon test** | | | | | |
|  | Condition | | Raw p-value | FDR-corrected p-value | | |
|  | Awake vs. N2 | | 0.0148 | 0.0222 | | |
|  | N1 vs. N2 | | 0.0101 | 0.0222 | | |
|  | Awake vs. N1 | | 0.1943 | 0.1943 | | |
| **Metastability** | **Friedman’s ANOVA** | | | | | |
|  | Source | SS | df | MS | Chi-sq | Prob>Chi-sq |
|  | Columns | 9.7 | 2 | 4.85 | 9.7 | 0.0078 |
|  | Error | 30.3 | 38 | 0.79737 |  |  |
|  | Total | 40 | 59 |  |  |  |
|  | **Post-hoc Wilcoxon test** | | | | | |
|  | Condition | | Raw p-value | FDR-corrected p-value | | |
|  | Awake vs. N2 | | 0.0111 | 0.0334 | | |
|  | N1 vs. N2 | | 0.1560 | 0.1560 | | |
|  | Awake vs. N1 | | 0.0438 | 0.0657 | | |
| **Complexity** | **Friedman’s ANOVA** | | | | | |
|  | Source | SS | df | MS | Chi-sq | Prob>Chi-sq |
|  | Columns | 6.1 | 2 | 3.05 | 6.1 | 0.0474 |
|  | Error | 33.9 | 38 | 0.89211 |  |  |
|  | Total | 40 | 59 |  |  |  |
|  | **Post-hoc Wilcoxon test** | | | | | |
|  | Condition | | Raw p-value | | FDR-corrected p-value | |
|  | Awake vs. N2 | | 0.0187 | | 0.0414 | |
|  | N1 vs. N2 | | 0.0276 | | 0.6274 | |
|  | Awake vs. N1 | | 0.6274 | | 0.0414 | |
